# Assessment of Molecular Mechanics-based Zn^2+^ Models in Mono- and Bimetallic Ligand Binding Sites

**DOI:** 10.1101/2021.06.28.450184

**Authors:** Okke Melse, Iris Antes

**Affiliations:** TUM Center for Functional Protein Assemblies and TUM School of Life Sciences, Technische Universität München, Emil-Erlenmeyer-Forum 8, 85354 Freising, Germany

## Abstract

Zn^2+^ ions play an important role in biology, but accurate sampling of metalloproteins using Molecular Mechanics remains challenging. Several models have been proposed to describe Zn^2+^ in biomolecular simulations, ranging from nonbonded models, employing classical 12-6 Lennard-Jones (LJ) potentials or extended LJ-potentials, to dummy-atom models and bonded models. We evaluated the performance of a large variety of these Zn^2+^ models in two challenging environments for which little is known about the performance of these methods, namely in a monometallic (Carbonic Anhydrase II) and a bimetallic ligand binding site (metallo-β-lactamase VIM-2). We focused on properties which are important for a stable, correct binding site description during molecular dynamics (MD) simulations, because a proper treatment of the metal coordination and forces are here essential. We observed that the strongest difference in performance of these Zn^2+^ models can be found in the description of interactions between Zn^2+^ and non-charged ligating atoms, such as the imidazole nitrogen in histidine residues. We further show that the nonbonded (12-6 LJ) models struggle most in the description of Zn^2+^-biomolecule interactions, while the inclusion of ion-induced dipole effects strongly improves the description between Zn^2+^ and non-charged ligating atoms. The octahedral dummy-atom models result in highly stable simulations and correct Zn^2+^ coordination, and are therefore highly suitable for binding sites containing an octahedral coordinated Zn^2+^ ion. The results from this evaluation study in ligand binding sites can guide structural studies of Zn^2+^ containing proteins, such as MD-refinement of docked ligand poses and long-term MD simulations.

## Introduction

Metalloproteins play an important role in many biological processes, making them interesting targets for drug design, but they are also highly suitable for many biotechnological applications.^1–4^ Furthermore, transition metals are of a complex nature, which increases the relevance of metalloproteins as biocatalysts for a large range of reactions. Understanding the ligand binding process in metal-containing binding sites is therefore key during the development of biocatalysts or during drug design. However, the complex chemistry of transition metal ions makes it challenging to properly describe the effect a metal ion has on its (biological) environment, and the interactions with the ligating atoms.^3^

Zn^2+^ is one of the most versatile biologically available metal ions, and often found in protein binding sites. For example, the recently presented ZincBind database^5^ counts 37,878 Zn^2+^-containing binding sites at the moment of writing. Zn^2+^ can play multiple roles in proteins, where one of the major roles is structural: stabilizing or enforcing a certain protein conformation, either to support the folding process or to bring the protein in an active conformation. Zinc’s ability to adopt multiple coordination numbers and geometries, typically 4, 5 and 6-coordinate Zn^2+^ complexes with tetrahedral/square planar, trigonal bipyramidal and octahedral coordination geometries are found in proteins, and the low energy barriers to switch between them makes Zn^2+^ an ideal metal ion to fulfill a variety of roles in biocatalysis.^6–9^

In classical molecular mechanics, the interaction between the metal ion and its environment is solely described via the nonbonded Coulomb and Lennard-Jones (LJ) potentials. Li *et al.*^10^ showed the difficulty of reproducing multiple experimentally measurable properties (here hydration free energy and ion-oxygen distance) with a single LJ-parameter set for divalent metal ions. Therefore, they developed three parameter sets, one optimized to reproduce hydration free energies, one for the ion-oxygen distance, and one reproducing a compromise of both. Subsequently, Li and Merz^11^ presented a modified LJ-potential, where they added a 1/r^4^ term to account for ion-induced dipole interactions. Applying this new potential, called the 12-6-4 LJ-type potential, one single parameter set for each metal ion could be developed which was able to reproduce both the hydration free energy and the ion-oxygen distance of the metal ion in water, as well as coordination numbers.^11–12^ Several studies have shown that this model can be used to predict a wide range of properties, often after some optimization of certain 12-6-4 LJ parameters. For example, Panteva *et al*.^13^ showed that a tuned 12-6-4 LJ-type model can be applied to accurately describe nucleic acid binding, and Sengupta *et al*.^14^ were able to describe chelate effects in an aqueous environment. More recently, Song *et al*.^15^ showed that this model can also reproduce the thermodynamics of transition metal ion binding in protein binding sites, as they were able to accurately calculate absolute binding free energies of Co^2+^ and Ni^2+^ in glyoxalase I, applying an optimized 12-6-4 LJ-type potential. Since this modification in the LJ-potential requires changes in the molecular dynamics (MD) code, only a limited amount of MD-engines support the use of this LJ-type. However, since of Amber18, support for this new LJ-potential has been implemented in the GPU-accelerated MD-code pmemd^16^, allowing long-term simulations applying this potential.

Another strategy to improve the description of the metal’s environment is the use of a bonded model.^17–18^ Here, the divalent coordinate bonds between the metal ion and its ligating atoms are modelled explicitly. However, this model generally requires system-specific parameterization of the newly introduced bonds and angles (dihedral angles are generally set to zero in this approach) as well as refitting partial charges of the metal ion(s) and the coordinating residues. Multiple strategies have been proposed to parameterize the bonded parameters. Thereof, the Seminario^19^ method is one of the most regularly applied methods, in which the force constants are derived from the Cartesian Hessian matrix. However, this method requires Quantum Mechanics (QM) calculations, which, besides the requirement for access to suitable QM-software and resources and thereby limiting the set-up of high-throughput calculations, may introduce problems. For example, the metal-complex could be unstable in the QM-geometry optimization, which makes the parameterization even more difficult. To avoid these drawbacks, Yu *et al*.^20^ recently presented the Extended Zinc Amber Force Field (EZAFF), which allows a fully empirical derivation of the bonded parameters required in the bonded model.

The major disadvantage of the bonded model is the lack of ligand exchange during the simulation. Since all metal ligands are explicitly bonded to the metal ion, a change of the metal environment cannot be sampled within this approach. Already in 1990, Åqvist and Warshel^21^ described a dummy-atom model (also called multisite model) for Mn^2+^, whereas in 1999, Pang^22^ was the first to parameterize this type of model for tetrahedral Zn^2+^. In this method, non-interacting dummy atoms carrying a portion of the metal’s mass and charge are placed around the metal ion, in a tetrahedral or octahedral geometry. The dummy-atom models can rotate freely around the metal ion without any energy penalty, but bonds between the metal ion and the dummy atoms keep them in the correct distance and geometry around the metal ion. Therefore, this alternative approach allows exchange of metal-ligating atoms, but still enforces/supports a certain coordination geometry due to the charge delocalization in the dummy atoms. More recently, Duarte *et al*.^23^ (re)parameterized the dummy-atom model for octahedral Zn^2+^ for the OPLS-AA force field, with the main focus to reproduce metal-oxygen distances, solvation free energy and coordination numbers. Two years later, Jiang *et al*.^24^ further refined the dummy-atom model parameters, using revised reference solvation free energies with Tissandier’s^25^ proton hydration free energies.

The presence of all those different models illustrates the complexity of a classical description of a metal ion. Since the nonbonded models, including the dummy-atom models, are parameterized in bulk water, there is little information about their performance in a protein environment. Yu *et al*.^20^ compared their EZAFF-derived bonded model in several Zn^2+^-containing systems with their bonded and nonbonded models and a selection of popular semi-empirical QM methods, based on geometry-optimized structures. However, in MD simulations, it is of great value to simulate around an equilibrium state, requiring analysis of the preferred conformational state of these Zn^2+^ models. Therefore, we performed long time scale MD simulations of the above mentioned models, analyzing the strengths and weaknesses of these Zn^2+^ models to reproduce experimental protein conformations around Zn^2+^ ions. This systematic evaluation was performed for Zn^2+^ ions in ligand binding sites, since proper modeling of the geometry of the zinc’s environment is most challenging in these cases, mostly because of the rather high amount of flexibility and presence of multiple potential Zn^2+^ ligating atoms. Additionally, a large amount of metalloproteins, such as metallo-β-lactamases, contain two Zn^2+^ ions in the binding site. Therefore, the performance of these Zn^2+^ models was evaluated in a monometallic as well as a bimetallic ligand binding site, using Carbonic Anhydrase II and the Verona integrin-encoded metallo-β-lactamase (VIM-2) as model systems.

## Materials and Methods

### Force Fields and Potential Function Forms

The nonbonded model in the AMBER force field is described in the following form:

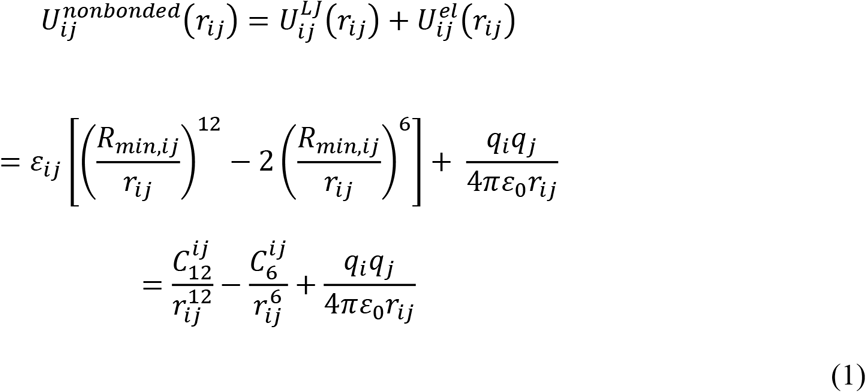

where r_ij_ is the distance between particle i and j, and R_min,ij_ and ɛ_ij_ are the distance between this particle pair at which the Lennard Jones (LJ) potential reaches its minimum and the LJ-well depth respectively, applying the Lorentz-Berthelot combining rules. q_i_ and q_j_ are the partial charges of the respective particles.

To improve the description of highly charged systems, Li and Merz^11^ presented a modified LJ potential for all interactions with metal ions, including a 1/r^4^ term describing the charge-induced dipole interactions. This potential is known as the 12-6-4 LJ-type potential because of its mathematical form:

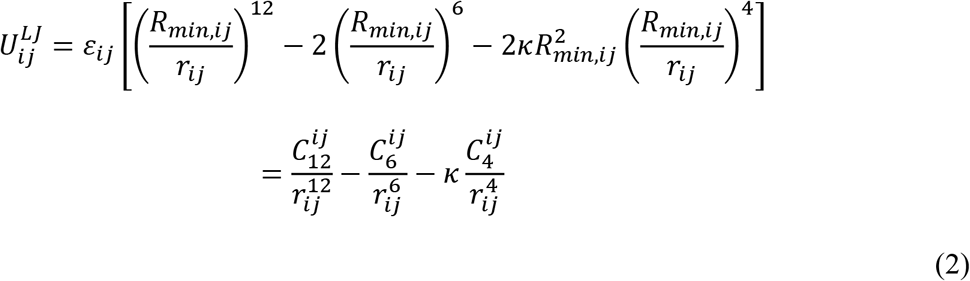

introducing κ as a scaling factor with the unit Å^−2^. 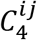 is defined as 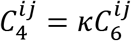 for interactions between the metal ion and the oxygen of water molecules, which is parameterized for a large set of divalent ions by Li and Merz.^11^ For the interactions of other atom types with metal ions, 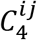is defined as follows:

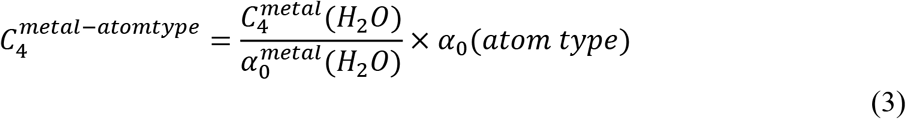

where α_0_ is the polarizability of the metal ion or respective atom type.

### Zn2+ Models

We evaluated four classical nonbonded (NB) parameter sets: the parameters from Merz^26^, which is the standard parameter set in the AMBER ff03 force field, as well as the three parameter sets from Li *et al*.^10^ The latter were optimized to reproduce the hydration free energy (NB-Li^HFE^), ion-oxygen distance (NB-Li^IOD^) or a compromise of both (NB-Li^CM^), where NB-Li^CM^ is the default parameter set in AMBER ff14SB. The major difference between these parameter sets lies in the value of the LJ well-depth, which ranges from 7.2·10^−4^ kcal·mol^−1^ to 1.5·10^−2^ kcal·mol^−1^. Because of the recent GPU implementation of the 12-6-4 LJ-type potential in the Amber MD-code, the 12-6-4 LJ-type parameter set could be evaluated in this study as well. Additionally, the three available Zn^2+^ dummy atom models (DU) were evaluated, containing one tetrahedral and two octahedral models. The dummy-atoms for the dummy-atom models were placed around the Zn^2+^ ions at a distance matching the equilibrium distance of the dummy-atom model applied. For the tetrahedral dummy atom model from Pang^27^, the revised force constant from Park *et al*.^28^ was used for the bonds involving dummy atoms. For the DU-Pang^mod^ model, the binding site was prepared according to the simulation setup from Pang^27–28^: metal-coordinating histidine residues were treated in their deprotonated form (histidine anion), and in the bimetallic binding site of VIM-2, Glu118 was protonated to form a hydrogen bond with the hydroxide ion. Additionally, all residues forming a hydrogen bond with a histidine anion were protonated. The parameters for the histidine anion and hydroxide ion were taken from Pang (2001)^27^.

Last, the performance of the bonded model (BM) was analyzed, with parameters derived from either the Seminario method or EZAFF. Since the partial charges used in this model are both retrieved from the same RESP procedure^29^, these values are identical between these two sets. MCPB.py^17^ was used to parameterize the divalent bonds for the bonded models: the positions of the hydrogen atoms from the large model generated by MCPB.py were optimized at the B3LYP/6-31G(d) level of theory. This optimized geometry was used for the RESP fitting, applying the ChgModB scheme, *i.e.* restraining all charges of the backbone heavy atoms to the values from the ff14SB force field, since this gave the best results in a study by Peters *et al*^18^. The van der Waals radius of the Zn^2+^ ion was set to 1.395 Å, taken from the IOD set of Li *et al*^10^. For the empirical model (BM-EZAFF), the bonded parameters were derived from the Extended Zinc Amber Force Field (EZAFF)^20^. For the Seminario model (BM-Seminario), a small model containing the Zn^2+^ ion and the coordination residues and ligand was generated with MCPB.py. The geometry of this small model was optimized at the B3LYP/6-31G(d) level of theory. The bonded parameters were derived applying the Seminario method on this geometry optimized small model. For VIM-2, the small model was initially optimized at the TPSS/def2-TZVP level of theory prior to the application of the Seminario method, to improve the description of the complex electronic structure in the bimetallic binding site. Additionally, a small van der Waals radius was placed on the hydrogen of the hydroxide ion, taken from the GAFF2 force field. All dihedrals involving a Zn^2+^ ion were set to zero, in agreement with the strategy applied in the MCPB procedure. The most relevant parameters of the evaluated Zn^2+^ models are shown in Table 1.

**Table 1.** Zn^2+^ Models Evaluated in this Study, with Parameters for the TIP3P Water Model.

| Parameter set | Model | Rmin/2 | $\epsilon$ | $\kappa$ | m<sub>zn</sub> | q<sub>zn</sub> | Reference |
| --- | --- | --- | --- | --- | --- | --- | --- |
| NB-Merz | Nonbonded LJ12-6 | 1.1000 | 0.012500 |  | 65.4 | +2 | Merz 1991<sup>26</sup> |
| NB-Li<sup>HFE</sup> | Nonbonded LJ12-6 (HFE set) | 1.1750 | 0.000716 |  | 65.4 | +2 | Li et al. 2013<sup>10</sup> |
| NB-Li<sup>IOD</sup> | Nonbonded LJ12-6 (IOD set) | 1.3950 | 0.014917 |  | 65.4 | +2 | Li et al. 2013<sup>10</sup> |
| NB-Li<sup>CM</sup> | Nonbonded LJ12-6 (CM set) | 1.2710 | 0.003303 |  | 65.4 | +2 | Li et al. 2013<sup>10</sup> |
| LJ1264 | 12-6-4 LJ-type | 1.4450 | 0.026628 | 1.623 | 65.4 | +2 | Li and Merz 2014<sup>11</sup> |
| DU-Pang<sup>a</sup> | Tetrahedral Dummy Atom Model | 3.1000 | 0.000001 |  | 53.83<br>(4x3.0) | 0<br>(4x+0.5) | Pang 2001<sup>27</sup> |
| DU-Duarte<sup>a</sup> | Octahedral Dummy Atom Model | 0.6814 | 112.734 |  | 47.39<br>(6x3.0) | -1<br>(6x+0.5) | Duarte et al. 2014<sup>23</sup> |
| DU-Jiang<sup>a</sup> | Octahedral Dummy Atom Model | 0.4895 | 100.7481 |  | 47.39<br>(6x3.0) | -1<br>(6x+0.5) | Jiang et al. 2016<sup>24</sup> |
| BM-Seminario<sup>b</sup> | Bonded model, Seminario | 1.3950 | 0.014917 |  | 65.4 | +0.5038/<br>+0.7351<sup>c</sup> | Li and Merz 2016<sup>17</sup> |
| BM-EZAFF<sup>b</sup> | Bonded model, Empirical (EZAFF) | 1.3950 | 0.014917 |  | 65.4 | +0.5038/<br>+0.7351<sup>c</sup> | Yu et al. 2018<sup>20</sup> |

### Simulation Setup

Carbonic Anhydrase II (CAII) in complex with the RA1 inhibitor (PDB ID: 5NXG) resolved at 1.2 Å resolution was used for the evaluation of Zn^2+^ models in monometallic systems, while the metallo-β-lactamase VIM-2 with the ANT-431 inhibitor (PDB ID: 6HF5), resolved at 1.8 Å resolution was applied as model system for the evaluation in bimetallic binding sites.^30–31^ During the ligand parameterization procedure, the ligands were protonated with Open Babel^32^ (v. 2.3.2.) at pH 7.0, keeping a deprotonated NH^-^ from the sulfonamide group of the CAII inhibitor, as this has been shown to be the state the ligand binds to the Zn^2+^ ion.^11, 32–33^ For the bonded and van der Waals parameters of the ligand atoms, the General Amber Force Field^34^ (GAFF) parameters were used. Charges of the ligand atoms for all but the bonded models were derived applying the RESP procedure based on a Merz-Singh-Kollman population analysis^35–36^ at a QM-geometry optimized structure of the inhibitor, which was performed at the HF/6-31G(d) level of theory, with Gaussian09 (revision E.01)^37^.

The protein-ligand complexes were prepared as follows: from all multi-resolved residues, the first occurrence was selected, and all water molecules within a sphere of 8 Å of the Zn^2+^ ion(s) were preserved. The protonation states of ionizable protein residues were predicted with the PROPKA3.0 software package^38–39^ at pH 7.0, followed by a visual check. For all standard protein residues, the AMBER ff14SB^40^ force field was applied. The system was solvated in a rectangular box containing TIP3P^41^ water, applying a buffer region of 12 Å around the protein atoms. The system was neutralized with Na^+^ ions using parameters from Joung and Cheatham^42^, using the LEaP module of the Amber18/AmberTools18 software package^16^. The bridging hydroxide ion in VIM-2 was parameterized as described in a previous study by Marion *et al.*^43^ containing a −1.3 e and +0.3 e partial charge for the oxygen and hydrogen atom respectively. The polarizability of the hydroxide oxygen, needed to calculate the C4 parameter in the 12-6-4 LJ-type model, was set to 2.03 Å^3^, taken from Shannon et al.^44^, while the polarizability of the hydrogen atom was set to zero, analogously to the hydrogen of water. The Zn^2+^ parameters from Table 1 were applied, and for the 12-6-4 LJ-type model, the C4 parameters were added with ParmEd.

### Simulation Procedure

An energy minimization was performed using sander from Amber18/AmberTools18 with the XMIN method (ntmin=3), thereby gradually adjusting the box size to bring the density from 0.8 g·cm^−3^ to 1.0 g·cm^−3^, in steps of 0.02 g·cm^−3^. During this minimization, a 20.0 kcal·mol·Å^−2^ positional restraint was applied to all protein atoms. When the target density was reached, an additional minimization was performed, applying the positional restraint solely to the binding site, defined as all residues with at least one atom in a sphere of 5 Å around the Zn^2+^ ion(s). During the heat-up procedure, the temperature was gradually increased to 300 K, while decreasing the positional restraint. The precise methodology is provided in the Supporting Information (Table S2 in the Supporting Information). The Langevin thermostat^45^ was applied with a collision frequency of 4.0 ps^−1^. During the MD simulations in the NPT ensemble, the Berendsen barostat^46^ was applied to keep the pressure at 1 bar, with a relaxation time of 1 ps and compressibility of 44.6·10^−6^·bar^−1^. Periodic boundary conditions were applied and the SHAKE algorithm^47^ was used to constrain all bonds involving a hydrogen atom at their equilibrium distance. A cut-off distance of 12 Å was used for all nonbonded interactions, while the particle mesh Ewald method^48^ was applied to describe long range electrostatics. The simulations were performed with an integration step size of 1 fs. After the heat-up, a 200 ns production MD simulation was performed at 300 K in a NPT ensemble, saving the atom coordinates and velocities every 10 ps. The heat-up and production simulations were conducted in three replicas, with the pmemd.cuda MD engine from Amber18.

### Trajectory analyses

From each replicon, the first 100 ns were considered as equilibration, thus the last 100 ns from the three replicas were merged, and analyzed as a single trajectory of 30000 frames. The distances and distance RMSDs between Zn^2+^ and the ligating atoms were calculated with cpptraj^49^. The coordination geometry was determined every 2 ns for each Zn^2+^ individually, using a python package FindGeo^50^. Every non-carbon heavy atom within 2.8 Å of the Zn^2+^ ion was considered as ligating atom, except the sulfur from the sulfonamide moiety in the CAII ligand.

## Results

We evaluated a large set of available Zn^2+^ models, which can be subdivided in four model types: classical nonbonded models (NB), the 12-6-4 LJ-type model (LJ1264), dummy-atom models (DU) and bonded models (BM). The performance of these methods was evaluated in the binding sites of CAII and VIM-2 (Figure 1), with the protonation state of the ligands and the binding site residues as supported by experiment (see Methods section). For the dummy atom model from Pang^27–28^, we conducted the performance evaluation with two different protonation states of the coordinating residues. The first evaluated setup (DU-Pang) is the protonation state identical as used for the other models while the second setup (DU-Pang^mod^) is simulated with deprotonated histidine residues and a protonated Asp118. The latter is the simulation setup equivalent to the setup used by Pang.

**Figure 1.**
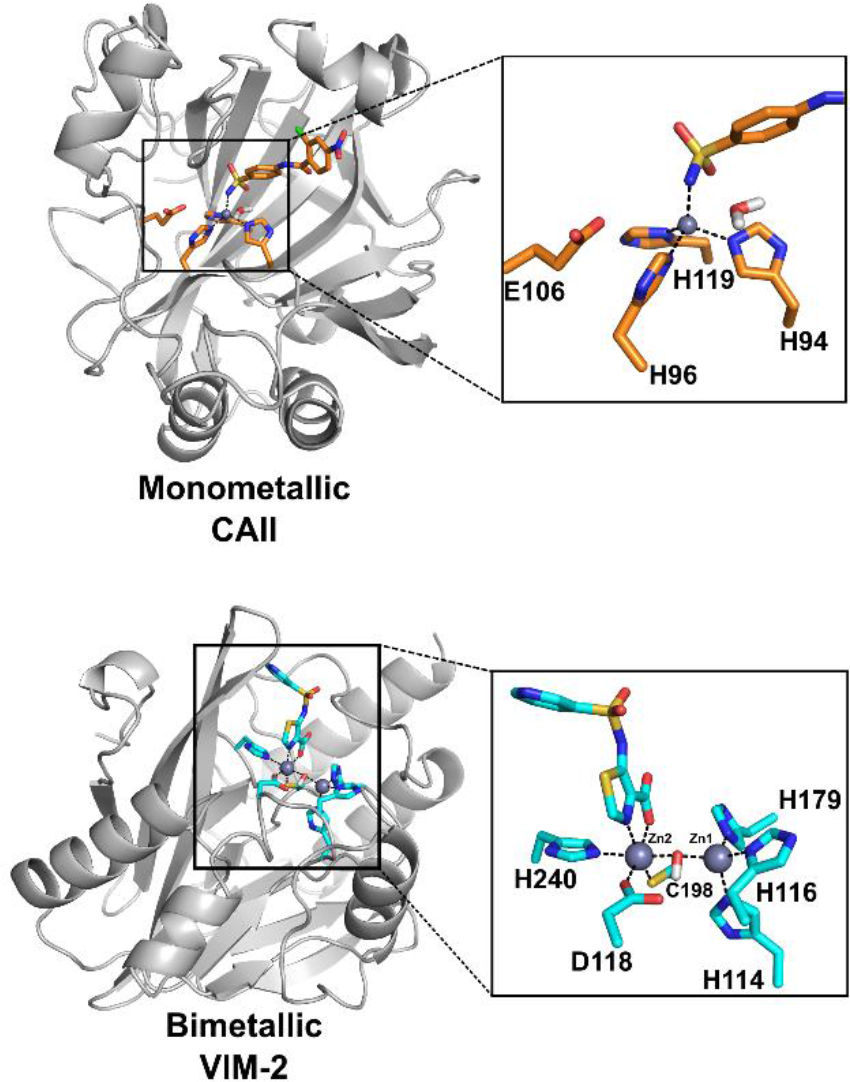
Evaluation systems used in this study. Carbonic Anhydrase II (CAII) was applied as model system for monometallic systems, while the β-lactamase VIM-2 was applied as model system for bimetallic systems. The proteins are shown in cartoon representations. The Zn^2+^ binding site, containing all coordinating residues and the inhibitor, as well as water molecules and protein residues which are close to the binding site and contain a heavy atom (potentially) able to coordinate the Zn^2+^ ion(s), are shown as licorice. The Zn^2+^ ions are illustrated as grey spheres. Dotted lines represent coordinate bonds.

To analyze the integrity of the Zn^2+^ binding site, the Root-Mean-Square Deviation was calculated over the interatomic distances (dRMSD) between the Zn^2+^ ion and a selection of atoms defining the structure of the binding site (Figure 2). The dRMSD is calculated with respect to the average distance (over all three replicas) simulated between the respective atom pair. Small dRMSD values thus indicates a stable ligation of the Zn^2+^ ion, because the interatomic distance between the Zn^2+^ ion and a ligating atom only slightly deviates from the average distance observed in the respective simulations. Large dRMSD values on the contrary indicate unstable ligation of the Zn^2+^ ion. The simulated Zn^2+^ binding sites by the nonbonded models show the largest distortion of the binding sites of all evaluated Zn^2+^ models, with the most unstable MD trajectories for NB-Merz and NB-Li^CM^. Besides the high average dRMSD values in the nonbonded models, the large error bars, especially for measures including histidine residues, illustrate that no stable conformation could be sampled within all replicate simulations. Addition of the ion-induced dipole effects via the 12-6-4 LJ-type potential improves the reproduction of the binding site significantly, especially for the Zn^2+^-His coordination bonds. Regarding the DU-Pang models, only simulations applying DU-Pang^mod^ show stable tetrahedral Zn^2+^ ions (CAII, and Zn_1_^2+^ in VIM-2). Remarkable is the strong performance of DU-Duarte and DU-Jiang, as these dummy-atom models simulate the VIM-2 binding site even closer to the experimental structure as the bonded models, illustrated by the dRMSD < 0.5 Å for all Zn^2+^-ligating atom distances. These dummy-atom models did not only reproduce the binding site for the octahedral coordinated Zn^2+^ ions, but also for the binding site around tetrahedral Zn^2+^.

**Figure 2.**
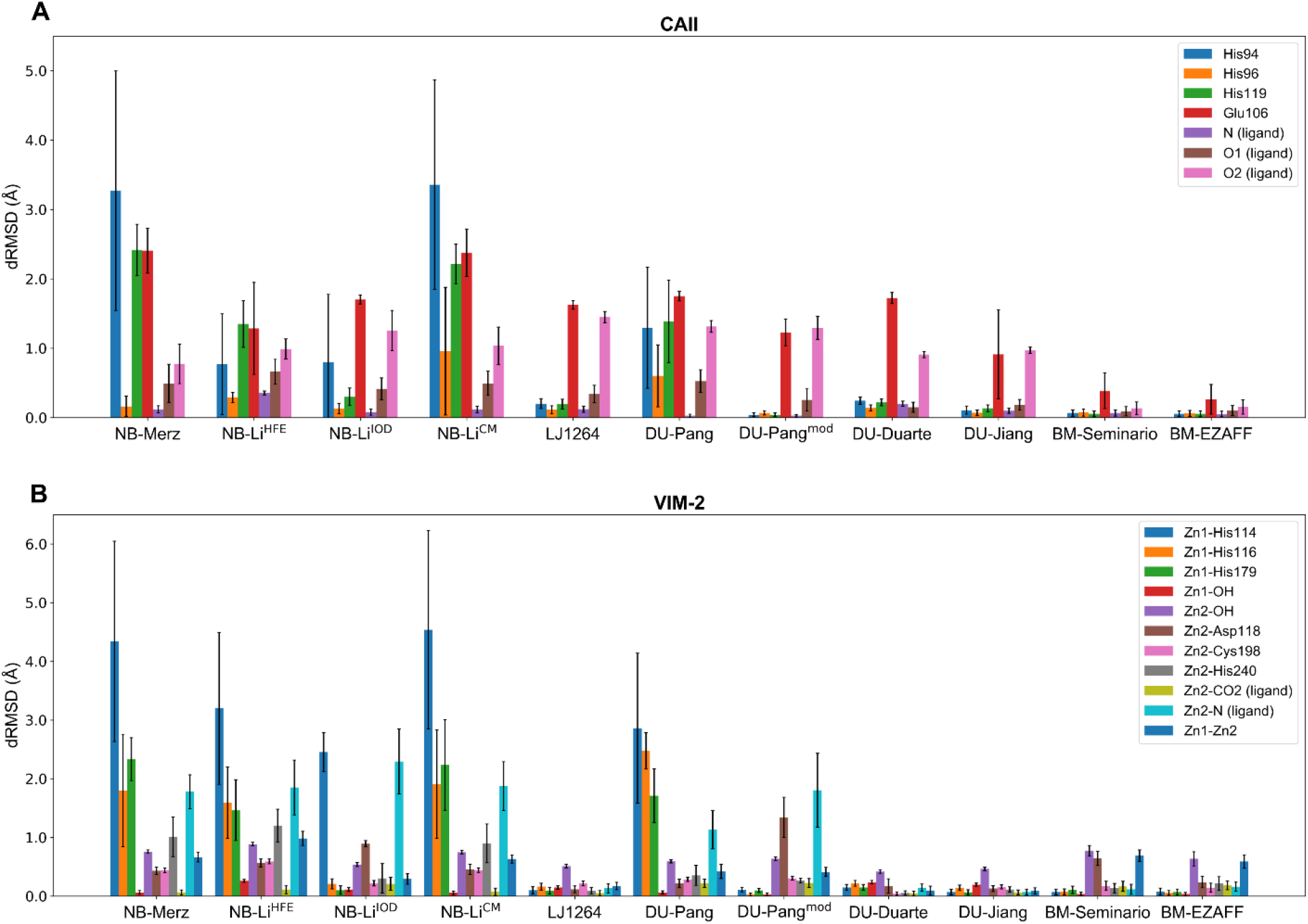
RMSD of interatomic distances (dRMSD) between Zn^2+^ and selected binding site atoms with respect to the average simulated interatomic distance of the respective atom pair, as a measure for the stability of the sampled Zn^2+^ binding site. The distances to histidine residues are defined as the distance between the respective Zn^2+^ ion to the ligating nitrogen of the histidine. The distances to an aspartate or glutamate residue are measured between the respective Zn^2+^ ion and the carboxyl carbon, since these side chains can rotate freely and contain two chemically equivalent oxygen atoms. A: distance RMSD in CAII. O1 (ligand) and O2 (ligand) represent the sulfonamide oxygen atoms, which are chemically equivalent, but an exchange of their position will lead to large changes in the binding pose of the ligand. N (ligand) represents the amine nitrogen of the sulfonamide moiety of the ligand. B: distance RMSD in VIM-2. The Zn_2_^2+^-CO2 (ligand) distance represents the distance between Zn_2_^2+^ and the carboxyl carbon from the inhibitor, where Zn_2_^2+^-N (ligand) is measured towards the nitrogen of the ligand’s thiazole moiety

Another important factor is the coordination geometry sampled by the Zn^2+^ models. This coordination geometry determines the exact position of the Zn^2+^ ligating atoms, which plays an important role in either stabilization of the protein structure, or introducing strain on certain bonds, for example to catalyze a chemical reaction. Therefore, we followed the coordination geometry of the Zn^2+^ ion(s) sampled during the simulations by determining the coordination geometry every 2 ns simulation time. The coordination geometry of the Zn^2+^ ion observed in the X-Ray structure CAII is tetrahedral, but this is only the most sampled coordination geometry by NB-Li^HFE^, DU-Pang^mod^, and the bonded models (Table 2). Almost all remaining Zn^2+^ models prefer an octahedral Zn^2+^ geometry, with 83% to 96% occurrence. The trigonal bipyramidal and square pyramidal geometries were barely sampled.

**Table 2.** Zn^2+^ Coordination Geometries Sampled in Monometallic CAII, in Percentage.

| Coordination geometry | NB-Merz | NB-$\text{Li}^{\text{HFE}}$ | NB-$\text{Li}^{\text{IOD}}$ | NB-$\text{Li}^{\text{CM}}$ | LJ1264 | DU-Pang | DU-Pang<sup>mod</sup> | DU-Duarte | DU-Jiang | BM-Seminario | BM-EZAFF |
| --- | --- | --- | --- | --- | --- | --- | --- | --- | --- | --- | --- |
| Tetrahedral<sup>a</sup> | 1.3 | 99.3 | 0.0 | 0.7 | 0.0 | 20.7 | 96.7 | 0.0 | 0.0 | 80.7 | 84.0 |
| Octahedral | 86.7 | 0.0 | 94.7 | 96.0 | 95.3 | 18.7 | 0.0 | 98.0 | 82.7 | 0.0 | 0.0 |
| Trigonal bipyramidal | 0.0 | 0.0 | 0.0 | 2.0 | 0.0 | 10.0 | 1.3 | 0.0 | 0.0 | 0.0 | 0.0 |
| Square pyramidal | 4.0 | 0.0 | 1.3 | 1.3 | 0.0 | 4.0 | 0.0 | 0.0 | 0.0 | 0.0 | 0.0 |
| Irregular geometry | 6.7 | 0.7 | 0.7 | 0.0 | 0.0 | 38.0 | 2.0 | 0.0 | 0.0 | 19.3 | 16.0 |
| Other<sup>b</sup> | 1.3 | 0.0 | 3.3 | 0.0 | 4.7 | 8.7 | 0.0 | 2.0 | 17.3 | 0.0 | 0.0 |

<sup>a</sup> Native coordination geometry, as observed in the X-Ray structure for CAII (PDB ID: 5NXG). <sup>b</sup> All coordination geometries which were visited less than 5 % in all evaluated $\text{Zn}^{2+}$ models in both CAII and VIM-2 were merged in this category. A complete list of all sampled coordination geometries for CAII is provided in the Supporting Information (Table S3).

A similar trend can be observed in the bimetallic system, VIM-2 (Table 3). The octahedral coordination geometry for Zn_1_^2+^ is clearly preferred by most nonbonded models, while a tetrahedral coordination geometry is found in the X-Ray structure of VIM-2. Besides the bonded models, only with the NB-Li^HFE^ and DU-Pang^mod^ models a tetrahedral coordination is sampled predominantly for Zn12+. Strikingly, the preferred coordination geometry for Zn22+, which is octahedral according to the X-Ray structure, was not always sampled in this octahedral geometry. With the NB-Merz, NB-Li^HFE^, NB-Li^CM^, and DU-Pang^mod^ models Zn_2_^2+^ is sampled mostly in a tetrahedral geometry. NB-Li^IOD^ and the 12-6-4 LJ-type model, however, simulated Zn_2_^2+^ in an octahedral geometry.

**Table 3.** Zn^2+^ Coordination Geometries Sampled in the Bimetallic System VIM-2, in Percentage

| Coordination geometry | NB-Merz | NB-Li<sup>HFE</sup> | NB-Li<sup>IOD</sup> | NB-Li<sup>CM</sup> | LJ1264 | DU-Pang | DU-Pang<sup>mod</sup> | DU-Duarte | DU-Jiang | BM-Seminario | BM-EZAFF |
| --- | --- | --- | --- | --- | --- | --- | --- | --- | --- | --- | --- |
| $\text{Zn}_1^{2+}$ geometry occupancy (%) | | | | | | | | | | | |
| Tetrahedral<sup>a</sup> | 0.0 | 96.0<sup>c</sup> | 0.0 | 0.0 | 0.0 | 4.7 | 99.3 | 0.0 | 0.0 | 89.3 | 94.0 |
| Octahedral | 95.3 | 0.0 | 100.0 | 96.7 | 100.0 | 22.7 | 0.0 | 100.0 | 100.0 | 0.0 | 0.0 |
| Trigonal bipyramidal | 0.0 | 1.3 | 0.0 | 0.0 | 0.0 | 40.7 | 0.0 | 0.0 | 0.0 | 0.0 | 0.7 |
| Square pyramidal | 4.0 | 0.0 | 0.0 | 2.7 | 0.0 | 0.0 | 0.0 | 0.0 | 0.0 | 0.0 | 0.0 |
| Irregular geometry | 0.7 | 2.7 | 0.0 | 0.7 | 0.0 | 27.3 | 0.0 | 0.0 | 0.0 | 0.7 | 4.7 |
| Other<sup>b</sup> | 0.0 | 0.0 | 0.0 | 0.0 | 0.0 | 4.7 | 0.7 | 0.0 | 0.0 | 10.0 | 0.7 |
| $\text{Zn}_2^{2+}$ geometry occupancy (%) | | | | | | | | | | | |
| Tetrahedral | 98.0 | 100.0 | 0.0 | 89.3 | 0.0 | 22.0 | 92.0 | 0.0 | 0.0 | 0.0 | 0.0 |
| Octahedral<sup>a</sup> | 0.0 | 0.0 | 43.3 | 0.0 | 99.3 | 0.7 | 0.0 | 100.0 | 100.0 | 100.0 | 100.0 |
| Trigonal bipyramidal | 0.7 | 0.0 | 0.0 | 2.0 | 0.0 | 17.3 | 0.0 | 0.0 | 0.0 | 0.0 | 0.0 |
| Square pyramidal | 0.0 | 0.0 | 0.0 | 0.0 | 0.0 | 0.0 | 0.0 | 0.0 | 0.0 | 0.0 | 0.0 |
| Irregular geometry | 1.3 | 0.0 | 18.7 | 8.7 | 0.0 | 58.0 | 8.0 | 0.0 | 0.0 | 0.0 | 0.0 |
| Other<sup>b</sup> | 0.0 | 0.0 | 38.0 | 0.0 | 0.7 | 2.0 | 0.0 | 0.0 | 0.0 | 0.0 | 0.0 |

<sup>a</sup> Native coordination geometry, as observed in the X-Ray structure for VIM-2 (PDB ID: 6HF5). <sup>b</sup> All coordination geometries which were visited less than 5 % in all evaluated $\text{Zn}^{2+}$ models in both CAII and VIM-2 were merged in this category. A complete list of all sampled coordination geometries for VIM-2 is provided in the Supporting Information (Table S4). <sup>c</sup> All coordinating ligands were water molecules or a hydroxide ion, and in one replicon the carboxylate oxygen of the ligand.

We also analyzed the preference of the Zn^2+^ models for certain ligating residues during the simulations (Figure 3). We defined a residue as ligating whenever their ligating atom is present within the coordination sphere of the Zn^2+^ ion, *i.e.* a sphere with a 2.8 Å radius around the Zn^2+^ ion. A normalized/average coordination number (CN) was calculated, where CN = 1 indicates a stable monodentate coordination between this residue and the Zn^2+^ ion. CN < 1 indicates a less stable interaction, since the respective coordination interaction was not constantly present over the entire simulation. CN values > 1 for a protein residue indicates that on average more than one ligating atom contributes to the Zn^2+^ coordination, which could occur for bidentate residues having more than one atom able to ligate the Zn^2+^ ion (*e.g.* Asp and Glu). CN > 1 for water molecules indicate that multiple water molecules coordinate the Zn^2+^ ion. This analysis shows a clear difference in the preference of the evaluated Zn^2+^ models towards specific ligating atoms. For example, the interaction with histidine residues is often not properly sampled, because the coordination is taken over by a charged residue or a water molecule. This evaluation gave further insights in the preference of some Zn^2+^ models for charged ligating atoms, or hard or soft bases, which will be discussed below.

**Figure 3.**
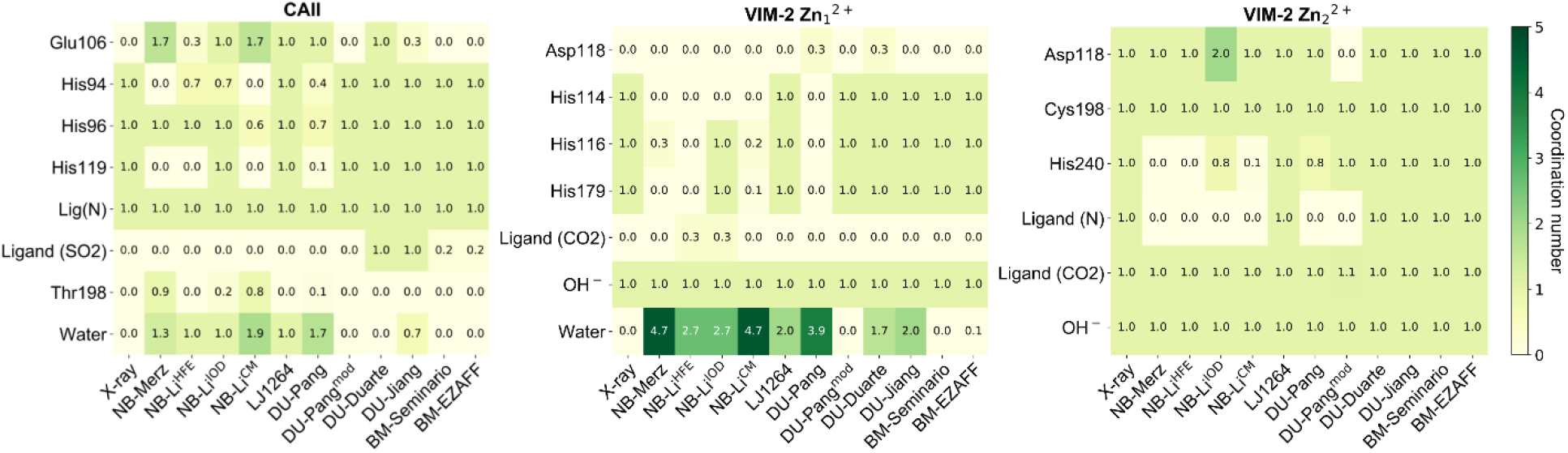
Contribution of ligating residues to the coordination of the Zn^2+^ ion during the simulations with the corresponding Zn^2+^ models. The values represent a normalized/average coordination number for each ligating residue. A residue is defined as ligating if the ligating atom of this residue is present within the coordination sphere (2.8 Å radius) around the Zn^2+^ ion. The heat maps are colored from light to dark green, ranging from small to large coordination numbers.

## Discussion

The performance of the Zn^2+^ models was analyzed in the binding sites of Carbonic anhydrase II and β-lactamase VIM-2, as model systems for monometallic and bimetallic binding sites respectively (Figure 1). CAII is very well suitable for the analysis of monometallic Zn^2+^ binding sites, since the Zn^2+^ ion in CAII is coordinated in a tetrahedral geometry, which is the dominant coordination geometry of Zn^2+^ monometallic binding sites.^8^ The Zn^2+^ ion is coordinated by three histidine residues and the nitrogen of the sulfonamide moiety of the inhibitor, again typical ligating residues for Zn^2+^. Due to the opposite charge between Glu106 and the Zn^2+^ ion, we hypothesized that certain Zn^2+^ models may overestimate this electrostatic interaction, and thereby falsely consider Glu106 as a coordinating residue. Therefore, the non-coordinating glutamate residue Glu106 close to the Zn^2+^ ion (4.0 Å and 5.5 Å between Zn^2+^ and both glutamate oxygen atoms respectively) was included in the analysis as well. Furthermore, Glu106 plays an important role in an elaborate hydrogen-bonding network in CAII: this glutamate orients binding site residues such that they can interact with the ligand, an effect which has been described in a variety of studies, illustrating the importance of proper sampling of this residue.^51–53^ Glu106 interacts with hydrogen bonds via a water molecule to Tyr7, and directly to the side chain of Thr199, which in turn interacts both via the backbone nitrogen and the side chain hydroxyl group to the sulfonamide moiety of the inhibitors.

The binding site of VIM-2 contains two Zn^2+^ ions, bridged by a hydroxide ion.^54–56^ Zn_1_^2+^ is coordinated in a tetrahedral geometry by three histidine residues and the bridging hydroxide, while Zn_2_^2+^ adopts an octahedral coordination geometry via an aspartate, cysteine and histidine residue, as well as the bridging hydroxide, the thiazole nitrogen and carboxylate oxygen of the inhibitor. Besides the interaction of the ligand with the Zn^2+^ ion, the ligand forms several hydrogen bonds with the protein: Arg228 forms a salt-bridge with the carboxylate group of the ligand, and Tyr67 forms an aromatic interaction with the ligand’s pyridine moiety.^31^ The two differently coordinated Zn^2+^ ions make this an interesting model system, allowing evaluation of the Zn^2+^ models to discriminate between different coordination geometries. Furthermore, the effect of the presence of a second Zn^2+^ ion within a single system on the Zn^2+^ model’s performance can be studied with this system.

In this study, we evaluated the performance of a variety of Zn^2+^ models in these two ligand binding sites, more precisely, in their ability to describe the structure of the Zn^2+^ coordination environment properly during MD simulations. We performed this study in ligand binding sites, since we think this is one of the most challenging case for the Zn^2+^ models, mainly due to the commonly rather high flexibility of ligand binding sites. Furthermore, ligand binding sites often contain a high number of solvent molecules, which are all able to coordinate Zn^2+^. Another reason to evaluate the performance in ligand binding sites is that proper structural sampling is highly important in the ligand binding site for a variety of practical applications, like (MD-based) molecular docking simulations and refinements, or studies with special focus on resolving chemical reaction pathways.

### Classical nonbonded models

Proper sampling of the metal’s coordination geometry is very important, mainly because the ion’s ability to bind its environment in a certain coordination geometry is one of the most important properties of transition metal ions in proteins. However, this property is often neglected.^57–58^ We found that the coordination geometry is least accurately reproduced by the nonbonded models compared to the other models in this study (Table 2 and 3), and that often incorrect ligating atoms are found (Figure 3 and 5), implicating that the standard 12-6 LJ potential is insufficient for proper description of Zn^2+^ ions. This observation follows previous findings from a recent evaluation study of Mg^2+^ models by Zuo and Liu.^58^ At first sight, the NB-Li^HFE^ model seems to work well, since a tetrahedral coordination geometry was sampled for the tetrahedral Zn^2+^ ions (CAII, and Zn_1_^2+^ in VIM-2). However, this tetrahedral coordination was not sampled with the correct ligating residues: in all replicas, His119 was replaced by a ligating water molecule and the ligating role of His94 was taken over by Glu106 in one of the replicate simulations (Figure 3). This artificial solvation of the Zn^2+^ ion in CAII was also observed for the NB-Merz and NB-Li^CM^ models, for which most coordination bonds with the histidine residues were broken and replaced by water molecules. Simulations of VIM-2 applying these nonbonded models again showed that all coordination bonds between the Zn^2+^ ions and the histidine residues, as well as the ligand’s thiazole nitrogen were broken and replaced by oxygen atoms, mostly from water molecules (Figure 4). Our data reveals a preference of the nonbonded models, with the exception of NB-Li^IOD^, towards hard bases as ligating atoms (as in water). The only exception is the interaction between Zn_2_^2+^ and Cys198 in VIM-2. This interaction remained intact in all Zn^2+^ models, despite the rather soft base character of cysteine. However, the interaction between Cys198 and Zn_2_^2+^ may be prevailed by the coulomb term here, because this residue was modelled in the deprotonated state, which may explain the observed stable interaction. We think that this preference of hard bases over soft bases as ligating atoms by the nonbonded models may partly be a result of the parameterization procedure, which is traditionally performed for a single ion in bulk solvent (water).

**Figure 4.**
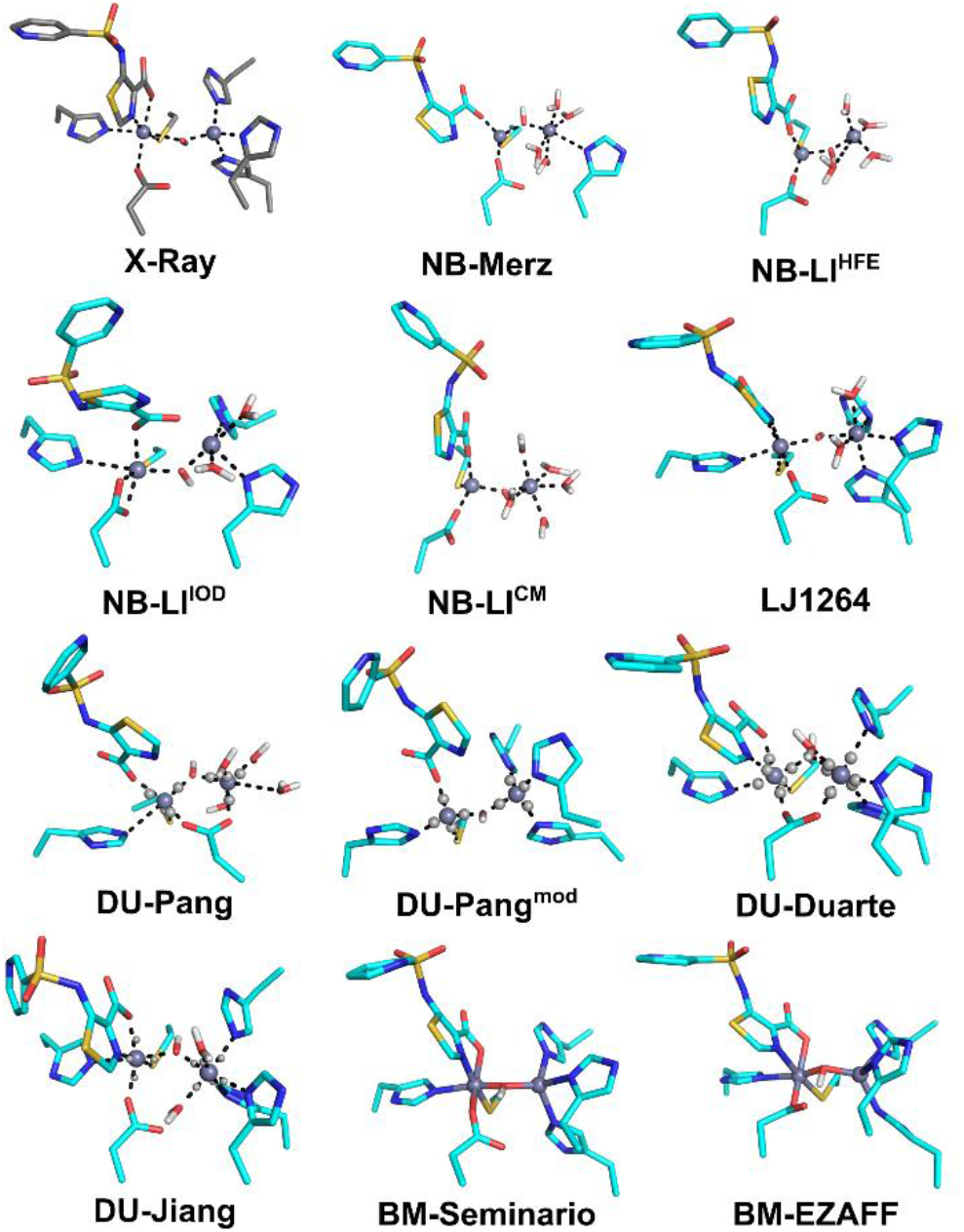
Sampled binding sites of VIM-2 by the evaluated Zn^2+^ models. Carbon atoms of the reference (X-Ray) and sampled structures are shown in respective grey and cyan. Zn^2+^ ions are represented as grey spheres, and the dummy-atoms as small light grey spheres. Dotted lines represent the coordination geometry.

**Figure 5.**
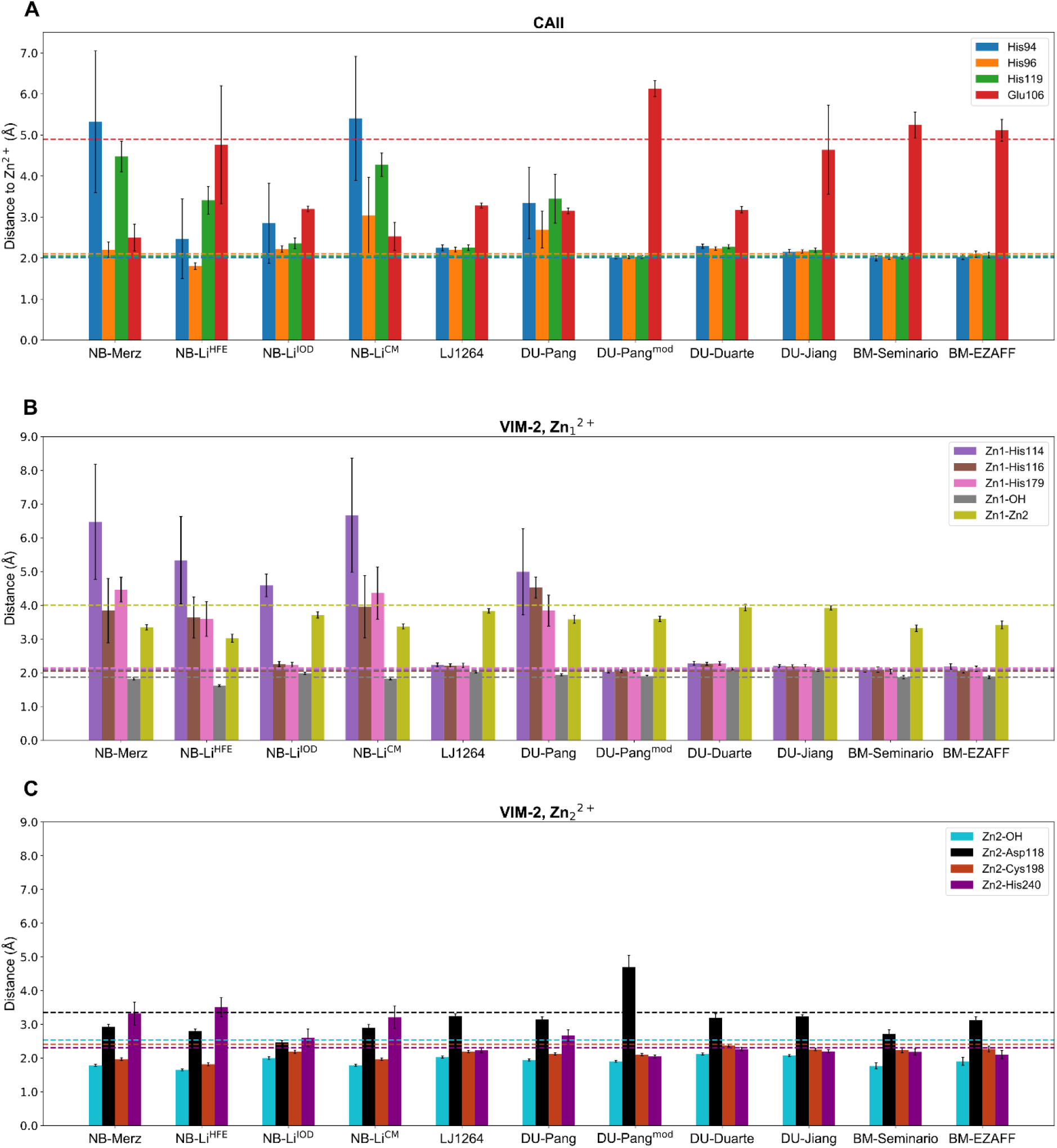
Interatomic distance between Zn^2+^ and selected binding site atoms, as a measure for binding site integrity for the Zn^2+^ ion in CAII (A), and Zn_1_^2+^ (B) and Zn_2_^2+^ (C) in VIM-2. The bars represent the average distance between the Zn^2+^ ion and the ligating residue during the simulation, while the dotted lines represent the value of the respective distance in the X-Ray structure. All distances to an aspartate or glutamate residue are measured to the side-chain’s carboxyl carbon, since both oxygen atoms are equivalent and can freely take over each-other’s function.

Both NB-Merz and NB-Li^CM^ models show a highly similar pattern in dRMSD measures and preferred coordinating residues (Figure 2, 5 and 6). Furthermore, both models preferred an octahedral coordination for CAII and Zn_1_^2+^ in VIM-2, while they simulated a tetrahedral coordination geometry for Zn_2_^2+^ (Table 2 and 3). The instable simulation trajectories and incorrect Zn^2+^ coordination sampled by these models result mainly from unstable binding of the Zn^2+^ ions, as the Zn^2+^ ions move up to 3 Å away from their crystallized position during the simulations (Figure S1 and S2 in the Supporting Information). NB-Li^IOD^, which was specifically optimized to reproduce ion-oxygen distances, properly described the Zn^2+^-His interactions for His96 and His119 in CAII and His116 and His179 in VIM-2, in contrast to the other 12-6 LJ nonbonded models as described above. However, this nonbonded model still failed to predict the interaction with His94 in CAII (in one replicon) and His114 in VIM-2 (in all replicas), as well as the thiazole nitrogen of the ligand in VIM-2, where the latter results in large deviations in the ligand binding pose (Figure 3 and 4). Additionally, an octahedral coordination geometry was sampled with the NB-Li^IOD^ model for all evaluated Zn^2+^ ions (Table 2 and 3), thus this model seems not to be suitable to reproduce tetrahedral binding sites. As described before, this preference for an octahedral coordination geometry could possibly be an artifact from the parameterization in water, since Zn^2+^ coordinates water in an octahedral geometry.^59–60^

**Figure 6.**
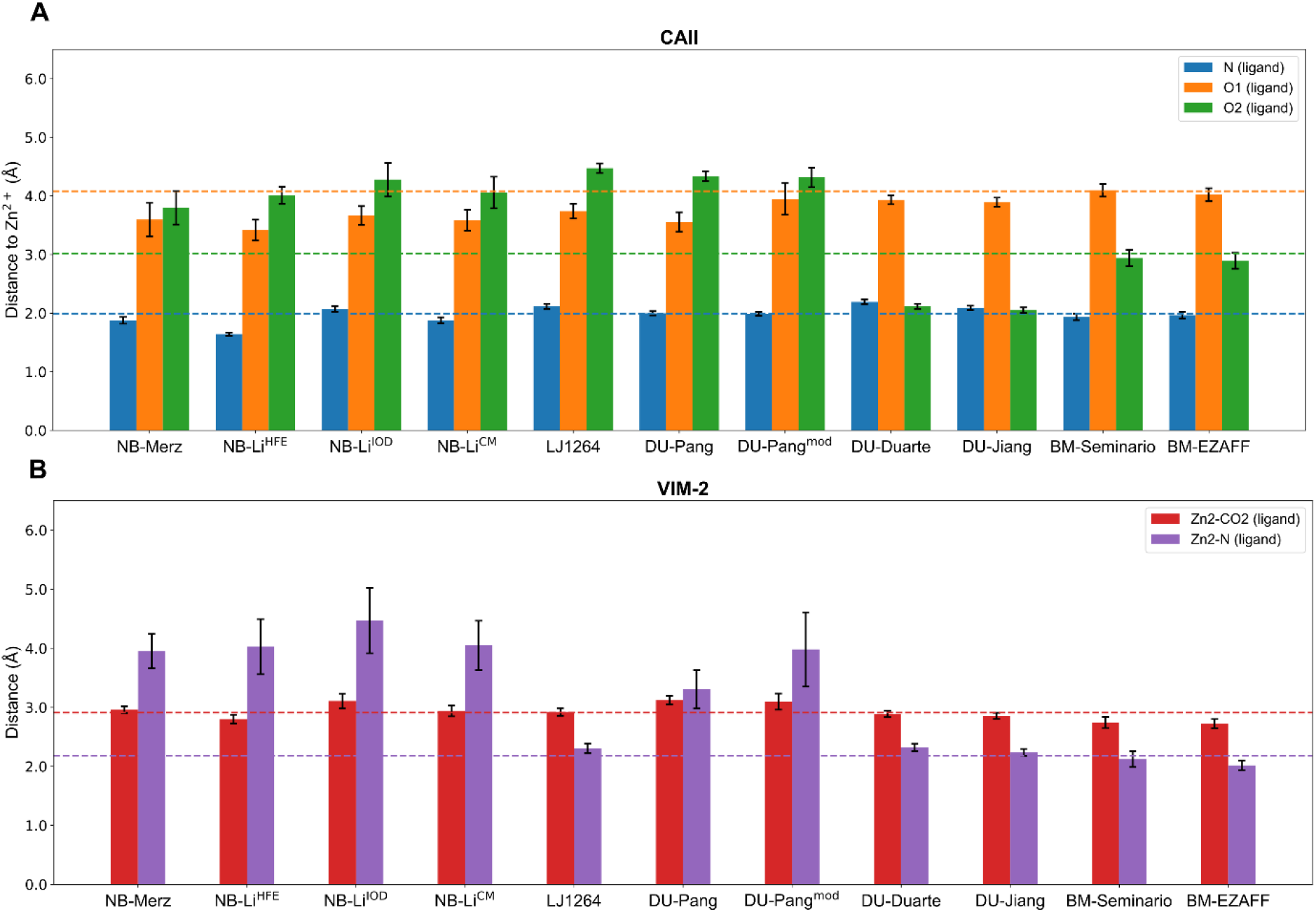
Interatomic distance between Zn^2+^ and electronegative atoms from the ligand in CAII (A) and VIM-2 (B). The bars represent the average distance between the Zn^2+^ ion and the ligating atom during the simulation, while the dotted lines represent the value of the respective distance in the X-Ray structure. Zn_2_^2+^-CO2 in VIM-2 is measured from the Zn^2+^ ion to the carboxyl carbon of the ligand, since both oxygen atoms are equivalent and can freely take over each-other’s function.

### Influence of ion-induced dipole effects

The idea from Li and Merz to include ion-induced dipole effects in the LJ-equation for all interactions containing a metal ion, *i.e.* the 12-6-4 LJ-type potential, leads to much better description of interactions between Zn^2+^ and particularly soft/borderline bases (Figure 3 and 5). However, we observed that the 12-6-4 LJ-type model almost solely models the Zn^2+^ ion in an octahedral coordination geometry (Table 2 and 3). This can be problematic whenever an accurate reproduction of the metal coordination is required, for example if MD simulations are used to refine docked ligand poses, because Zn^2+^ often has a tetrahedral coordination in protein structures.^8^ To accomplish this octahedral coordination geometry in the tetrahedral binding sites CAII, and Zn_1_^2+^ in VIM-2, additional water molecules were ligated in simulations applying the 12-6-4 LJ-type potential (Figure 3), an effect which was also observed in simulations with other metal ions.^61^ Furthermore, our simulations of CAII applying the 12-6-4 LJ-type model still overshoot the interaction between Zn^2+^ and the charged Glu106, as illustrated by the too short distance between them (Figure 5). Despite these inaccuracies, this modified LJ-potential leads to much better description of Zn^2+^ coordination bonds in a protein environment, and thereby increases the stability of the binding site geometries considerably compared to the classical 12-6 LJ nonbonded models (Figure 2). This better metal description also improved the sampling of the ligand binding pose in VIM-2 (Figure 4): the heavy-atom RMSD of the ligand is 1.8 Å in the simulations applying the 12-6-4 LJ-type potential, using the crystal structure as reference, while the ligand RMSD of the other nonbonded models are higher than 2.8 Å.

### Dummy atom models and the influence of protonation states

The simulation of the tetrahedral binding sites can be improved substantially by the DU-Pang^mod^ dummy atom model. This model sampled the tetrahedral binding sites almost exclusively in their correct coordination geometry: 96.7% and 99.3% for CAII, and Zn_1_^2+^ in VIM-2 respectively (Table 2 and 3). However, this result could only be reached applying the (artificial) deprotonated histidine residues and modified protonation states of the other coordinating residues, as used by Pang in his publication as well.^22^ The performance worsened drastically once the same protonation states were used as in all other Zn^2+^ models, *i.e.* neutral histidine residues, (named DU-Pang in this study): no stable coordination geometries could be found in these cases, only 20.7% and 4.7% of the simulation showed a tetrahedral coordination for CAII and Zn_1_^2+^ in VIM-2, respectively. Furthermore, following Pang’s rules to determine which residues should be protonated, the required protonation state of Asp118 in VIM-2 remains ambiguous. Asp118 should be deprotonated according to Pang’s rules because of its interaction with Zn22+, yet it must be protonated because of its hydrogen-bonding with the hydroxide ion. Therefore, we performed the simulations applying the DU-Pang^mod^ model with both protonated and deprotonated Asp118: the simulation containing a deprotonated Asp118 resulted in a highly distorted binding site, thereby losing ligand binding (data not shown), while the simulation with a protonated Asp118 stayed rather close to the experimentally observed structure. Therefore, the simulation with the protonated Asp118 represents the DU-Pang^mod^ model in this study. This protonated Asp118 did however not form a hydrogen bond with the hydroxide ion during the simulation, which was the rationalization to protonate this residue. In contrast, Asp118 formed a hydrogen bond with solvent molecules and thereby losing its coordination of the Zn^2+^ ion (Figure 3 and 5). Both DU-Pang and DU-Pang^mod^ were not able to reproduce the octahedral binding geometry from Zn_2_^2+^ in VIM-2. With the DU-Pang model an octahedral geometry was simulated in only 0.7% of the simulation time, and with the DU-Pang^mod^ model a tetrahedral geometry was found instead (Table 3). Thus, DU-Pang^mod^ performs very well for tetrahedral Zn^2+^ ions, but the performance is highly sensitive on the choice of protonation state of the coordinating residues and is not suitable to reproduce another coordination geometry than tetrahedral.

The other Zn^2+^ dummy atom models, DU-Duarte and DU-Jiang, the latter being a re-parameterization of DU-Duarte based on revised solvation free energies^24^, were both parameterized for octahedral Zn^2+^ ions, and therefore contain six dummy-atoms.^23–24^ Despite their coordination geometry specific parameterization, they also performed quite well for the tetrahedral Zn^2+^ ions in this study. The six dummy-atoms enforce an octahedral coordination geometry, also for the tetrahedral Zn^2+^ ions (Table 2 and 3), but this had only a minor effect on the positions and stability of the coordinating residues (Figure 5). The additional coordination partners required to fulfill the octahedral coordination geometry for Zn_1_^2+^ in VIM-2 were mainly water molecules, which we also observed for simulations applying the 12-6-4 LJ-type potential (Figure 3). In CAII, in both, the DU-Duarte and DU-Jiang models, a sulfonamide oxygen was used as an additional ligating atom, but this barely affected the ligand binding orientation. Using the DU-Duarte model, an additional ligation with Glu106 was observed in all replicas, yet this glutamate kept its important interaction with Thr199, which was shown to be a conserved interaction in CAII. With the DU-Jiang model, ligation of Glu106 was found only in one of the replicas, while a water molecule was used in the other two. Thus, this resulted in an almost perfect reproduction of the Zn^2+^ binding site and in very stable simulation trajectories (Figure 2), both for the monometallic and bimetallic system, despite the wrong coordination environment of the tetrahedral Zn^2+^ ions. Both models, DU-Duarte and DU-Jiang, perform very well for the octahedral Zn_2_^2+^ in VIM-2, applying the correct ligating residues over the entire simulation (Figure 3).

Finally, we also analyzed if a tetrahedral and octahedral dummy-atom model can be combined in the bimetallic system VIM-2. Therefore, we performed simulations modelling the tetrahedral Zn_1_^2+^ by either DU-Pang or DU-Pang^mod^, and the octahedral Zn_2_^2+^ by DU-Jiang. The simulations applying the combined DU-Pang^mod^ and DU-Jiang models resulted in highly stable simulations and reproduced the coordination geometry of both ions correctly (Figure S3 and Table S4 in Supporting Information). However, in the combined model simulations, one still needs to (artificially) deprotonate the residues around Zn_1_^2+^ in order to sample a correct coordination geometry. Simulations with protonated and deprotonated Asp118 were similar, whereas the first performed best in the reproduction of the metal environment.

### Restraining metal-coordination via explicit bonds

The bonded models reproduced the Zn^2+^ sites rather accurately, which could be expected because of the system-specific parameterization and application of explicit bonds between the Zn^2+^ ions and ligating atoms. Correct usage of the ligating residues, proper reproduction of the coordination geometry and highly stable simulation trajectories were observed. It is however surprising that simulations of VIM-2 applying the dummy-atom models DU-Duarte and DU-Jiang were even more stable than the bonded models, and the simulated bimetallic binding site geometries were closer to the experimental structure than for both bonded models, while not having the disadvantage of fixed ligating residues. Especially the distance between the two Zn^2+^ ions (which were connected via explicit bonds in the bonded model via the hydroxide ion) was simulated closer to the experimental value by these dummy-atom models, as shown in Figure 5. It needs to be noted here that BM-Seminario is parameterized based on a QM-geometry optimized structure, during which the distance between the Zn^2+^ ions reduced to 3.7 Å, with respect to 4.0 Å in the X-Ray structure. However, the distance between the two Zn^2+^ ions during the BM-Seminario simulations was even smaller (3.3 Å). Finally, we barely observed any performance differences between BM-Seminario and BM-EZAFF, indicating that the empirical derivation of the bonded parameters works well, both for monometallic and bimetallic binding sites.

## Conclusions

Based on our evaluation of a large variety of Zn^2+^ models, we show that the nonbonded models applying the standard 12-6 LJ potential are too limited to accurately describe the conformation of a flexible Zn^2+^-containing protein binding site. Especially interactions between the Zn^2+^ ion and non-charged ligating atoms are underestimated, such as the imidazole nitrogen in histidine residues. These ligating atoms are often replaced by water molecules or charged residues during the simulation. The performance can be improved by the introduction of the ion-induced dipole interactions via the 12-6-4 LJ-type potential, which thanks to the GPU-implementation in AMBER18 is now also applicable for long time scale simulations. However, we showed that the 12-6-4 LJ-type model strongly prefers an octahedral coordination geometry and overestimates interactions with charged atoms, but this had only a minor effect on the binding site conformations sampled in this study. The dummy-atom models are a good alternative to the nonbonded models, from which DU-Pang^mod^ reproduces a tetrahedral coordination geometry fairly well, but its performance depends strongly on the not always rational protonation state of the metal’s environment. Since many Zn^2+^ ions in proteins are found in a tetrahedral coordination geometry, the development of a tetrahedral dummy-atom model parameterized specifically in a protein environment would be welcome. The octahedral dummy-atom models DU-Duarte and DU-Jiang are good choices for Zn^2+^ ions coordinated in an octahedral geometry because of their solid performance in this study regarding the reproduction of the metal’s environment. We also observed that these models perform rather well for tetrahedral coordinated Zn^2+^, but still attract water molecules to fill the coordination sphere. The difference in performance between DU-Duarte and DU-Jiang is minor, whereas the latter showed the best performance in this study. These octahedral dummy-atom models perform very similar, in the case of the bimetallic VIM-2 binding site even slightly better than the bonded models in this study. However, the latter may presumably still be the best choice whenever a fixed coordination geometry needs to be sampled, especially if the required geometry is disturbed and does not match any of the parameterized geometries. The combination of different dummy models based on their coordination geometry in the bimetallic site led to very promising results, indicating that it would be very useful to provide parameter sets for these models for different coordination environments. Because of the almost identical performance of BM-Seminario and BM-EZAFF in this evaluation study, the EZAFF method to parameterize the bonded model will often suffice, having the advantage of being independent of QM-calculations for its parameterization, except for charge fitting.

## Supporting Information

The Supporting Information is available free of charge (PDF). Figure S1. Zinc displacement.

Figure S2. Sampled coordination geometries in CAII.

Figure S3. Performance of combining DU-Pang and DU-Jiang for the bimetallic system VIM-2. Table S1. Bonded parameters applied in bonded models.

Table S2. Overview of heat-up protocol applied in this study.

Table S3. Full list of sampled Zn^2+^ coordination geometries in CAII. Table S4. Full list of sampled Zn^2+^ coordination geometries in VIM-2.

## Acknowledgements

The authors thank the Deutsche Forschungsgemeinschaft for financial support via SFB 749, project C08 and CIPSM.

## References

1. Riccardi, L.; Genna, V.; De Vivo, M., Metal–ligand interactions in drug design. Nat. Rev. Chem. 2018, 2 (7), 100–112.

2. Parac-Vogt, T. N.; Erxleben, A.; Schenk, G.; Prabhakar, R., Editorial: Advances in the Development of Artificial Metalloenzymes. Front. Chem. 2019, 7 (599).

3. Li, P.; Merz, K. M., Metal Ion Modeling Using Classical Mechanics. Chem. Rev. 2017, 117 (3), 1564–1686.

4. Bornscheuer, U. T., The fourth wave of biocatalysis is approaching. Philos. Trans. R. Soc. A 2018, 376 (2110), 20170063.

5. Ireland, S. M.; Martin, A. C. R., ZincBind—the database of zinc binding sites. Database 2019, 2019.

6. Dudev, M.; Wang, J.; Dudev, T.; Lim, C., Factors Governing the Metal Coordination Number in Metal Complexes from Cambridge Structural Database Analyses. J. Phys. Chem. B 2006, 110 (4), 1889–1895.

7. Kuppuraj, G.; Dudev, M.; Lim, C., Factors Governing Metal−Ligand Distances and Coordination Geometries of Metal Complexes. J. Phys. Chem. B 2009, 113 (9), 2952–2960.

8. Laitaoja, M.; Valjakka, J.; Jänis, J., Zinc Coordination Spheres in Protein Structures. Inorganic Chemistry 2013, 52 (19), 10983–10991.

9. Vahrenkamp, H., Why does nature use zinc-a personal view. Dalton Trans. 2007, (42), 4751–4759.

10. Li, P.; Roberts, B. P.; Chakravorty, D. K.; Merz, K. M., Rational Design of Particle Mesh Ewald Compatible Lennard-Jones Parameters for +2 Metal Cations in Explicit Solvent. J. Chem. Theory Comput. 2013, 9 (6), 2733–2748.

11. Li, P.; Merz, K. M., Taking into Account the Ion-Induced Dipole Interaction in the Nonbonded Model of Ions. J. Chem. Theory Comput. 2014, 10 (1), 289–297.

12. Li, P.; Song, L. F.; Merz, K. M., Systematic Parameterization of Monovalent Ions Employing the Nonbonded Model. J. Chem. Theory Comput. 2015, 11 (4), 1645–1657.

13. Panteva, M. T.; Giambaşu, G. M.; York, D. M., Force Field for Mg2+, Mn2+, Zn2+, and Cd2+ Ions That Have Balanced Interactions with Nucleic Acids. J. Phys. Chem. B 2015, 119 (50), 15460–15470.

14. Sengupta, A.; Seitz, A.; Merz, K. M., Simulating the Chelate Effect. Journal of the American Chemical Society 2018, 140 (45), 15166–15169.

15. Song, L. F.; Sengupta, A.; Merz, K. M., Thermodynamics of Transition Metal Ion Binding to Proteins. Journal of the American Chemical Society 2020, 142 (13), 6365–6374.

16. Case, D. A.; Ben-Shalom, I. Y.; Brozell, S. R.; Cerutti, D. S.; Cheatham, T. E., III; Cruzeiro, V. W. D.; Darden, T. A.; Duke, R. E.; Ghoreishi, D.; Gilson, M. K.; Gohlke, H.; Goetz, A. W.; Greene, D.; Harris, R.; Homeyer, N.; Huang, Y.; Izadi, S.; Kovalenko, A.; Kurtzman, T.; Lee, T. S.; LeGrand, S.; Li, P.; Lin, C.; Liu, J.; Luchko, T.; Luo, R.; Mermelstein, D. J.; Merz, K. M.; Miao, Y.; Monard, G.; Nguyen, C.; Nguyen, H.; Omelyan, I.; Onufriev, A.; Pan, F.; Qi, R.; Roe, D. R.; Roitberg, A.; Sagui, C.; Schott-Verdugo, S.; Shen, J.; Simmerling, C. L.; Smith, J.; Salomon-Ferrer, R.; Swails, J.; Walker, R. C.; Wang, J.; Wei, H.; Wolf, R. M.; Wu, X.; Xiao, L.; York, D. M.; Kollman, P. A., AMBER 18. University of California, San Francisco: 2018.

17. Li, P.; Merz, K. M., MCPB.py: A Python Based Metal Center Parameter Builder. J. Chem. Inf. Model. 2016, 56 (4), 599–604.

18. Peters, M. B.; Yang, Y.; Wang, B.; Füsti-Molnár, L.; Weaver, M. N.; Merz, K. M., Structural Survey of Zinc-Containing Proteins and Development of the Zinc AMBER Force Field (ZAFF). J. Chem. Theory Comput. 2010, 6 (9), 2935–2947.

19. Seminario, J. M., Calculation of intramolecular force fields from second-derivative tensors. International Journal of Quantum Chemistry 1996, 60 (7), 1271–1277.

20. Yu, Z.; Li, P.; Merz, K. M., Extended Zinc AMBER Force Field (EZAFF). J. Chem. Theory Comput. 2018, 14 (1), 242–254.

21. Aaqvist, J.; Warshel, A., Free energy relationships in metalloenzyme-catalyzed reactions. Calculations of the effects of metal ion substitutions in staphylococcal nuclease. Journal of the American Chemical Society 1990, 112 (8), 2860–2868.

22. Pang, Y.-P., Novel Zinc Protein Molecular Dynamics Simulations: Steps Toward Antiangiogenesis for Cancer Treatment. J. Mol. Model. 1999, 5 (10), 196–202.

23. Duarte, F.; Bauer, P.; Barrozo, A.; Amrein, B. A.; Purg, M.; Åqvist, J.; Kamerlin, S. C. L., Force Field Independent Metal Parameters Using a Nonbonded Dummy Model. J. Phys. Chem. B 2014, 118 (16), 4351–4362.

24. Jiang, Y.; Zhang, H.; Tan, T., Rational Design of Methodology-Independent Metal Parameters Using a Nonbonded Dummy Model. J. Chem. Theory Comput. 2016, 12 (7), 3250–3260.

25. Tissandier, M. D.; Cowen, K. A.; Feng, W. Y.; Gundlach, E.; Cohen, M. H.; Earhart, A. D.; Coe, J. V.; Tuttle, T. R., The Proton’s Absolute Aqueous Enthalpy and Gibbs Free Energy of Solvation from Cluster-Ion Solvation Data. J. Phys. Chem. A 1998, 102 (40), 7787–7794.

26. Merz, K. M., Carbon dioxide binding to human carbonic anhydrase II. Journal of the American Chemical Society 1991, 113 (2), 406–411.

27. Pang, Y.-P., Successful molecular dynamics simulation of two zinc complexes bridged by a hydroxide in phosphotriesterase using the cationic dummy atom method. Proteins: Struct. Funct. Bioinform. 2001, 45 (3), 183–189.

28. Park, J. G.; Sill, P. C.; Makiyi, E. F.; Garcia-Sosa, A. T.; Millard, C. B.; Schmidt, J. J.; Pang, Y.-P., Serotype-selective, small-molecule inhibitors of the zinc endopeptidase of botulinum neurotoxin serotype A. Biorg. Med. Chem. 2006, 14 (2), 395–408.

29. Bayly, C. I.; Cieplak, P.; Cornell, W.; Kollman, P. A., A well-behaved electrostatic potential based method using charge restraints for deriving atomic charges: the RESP model. J. Phys. Chem. 1993, 97 (40), 10269–10280.

30. Pecina, A.; Brynda, J.; Vrzal, L.; Gnanasekaran, R.; Hořejší, M.; Eyrilmez, S. M.; Řezáč, J.; Lepšík, M.; Řezáčová, P.; Hobza, P.; Majer, P.; Veverka, V.; Fanfrlík, J., Ranking Power of the SQM/COSMO Scoring Function on Carbonic Anhydrase II–Inhibitor Complexes. ChemPhysChem 2018, 19 (7), 873–879.

31. Leiris, S.; Coelho, A.; Castandet, J.; Bayet, M.; Lozano, C.; Bougnon, J.; Bousquet, J.; Everett, M.; Lemonnier, M.; Sprynski, N.; Zalacain, M.; Pallin, T. D.; Cramp, M. C.; Jennings, N.; Raphy, G.; Jones, M. W.; Pattipati, R.; Shankar, B.; Sivasubrahmanyam, R.; Soodhagani, A. K.; Juventhala, R. R.; Pottabathini, N.; Pothukanuri, S.; Benvenuti, M.; Pozzi, C.; Mangani, S.; De Luca, F.; Cerboni, G.; Docquier, J.-D.; Davies, D. T., SAR Studies Leading to the Identification of a Novel Series of Metallo-β-lactamase Inhibitors for the Treatment of Carbapenem-Resistant Enterobacteriaceae Infections That Display Efficacy in an Animal Infection Model. ACS Infect. Dis. 2019, 5 (1), 131–140.

32. O’Boyle, N. M.; Banck, M.; James, C. A.; Morley, C.; Vandermeersch, T.; Hutchison, G. R., Open Babel: An open chemical toolbox. J. Cheminform. 2011, 3 (1), 33.

33. Pecina, A.; Lepšík, M.; Řezáč, J.; Brynda, J.; Mader, P.; Řezáčová, P.; Hobza, P.; Fanfrlík, J., QM/MM Calculations Reveal the Different Nature of the Interaction of Two Carborane-Based Sulfamide Inhibitors of Human Carbonic Anhydrase II. J. Phys. Chem. B 2013, 117 (50), 16096–16104.

34. Wang, J.; Wolf, R. M.; Caldwell, J. W.; Kollman, P. A.; Case, D. A., Development and testing of a general amber force field. J. Comput. Chem. 2004, 25 (9), 1157–74.

35. Besler, B. H.; Merz, K. M. Jr; Kollman, P. A., Atomic charges derived from semiempirical methods. J. Comput. Chem. 1990, 11 (4), 431–439.

36. Singh, U. C.; Kollman, P. A., An approach to computing electrostatic charges for molecules. J. Comput. Chem. 1984, 5 (2), 129–145.

37. Frisch, M. J.; Trucks, G. W.; Schlegel, H. B.; Scuseria, G. E.; Robb, M. A.; Cheeseman, J. R.; Scalmani, G.; Barone, V.; Mennucci, B.; Petersson, G. A.; Nakatsuji, H.; Caricato, M.; Li, X.; Hratchian, H. P.; Izmaylov, A. F.; Bloino, J.; Zheng, G.; Sonnenberg, J. L.; Hada, M.; Ehara, M.; Toyota, K.; Fukuda, R.; Hasegawa, J.; Ishida, M.; Nakajima, T.; Honda, Y.; Kitao, O.; Nakai, H.; Vreven, T.; Montgomery, J. A. J.; Peralta, J. E.; Ogliaro, F.; Bearpark, M.; Heyd, J. J.; Brothers, E.; Kudin, K. N.; Staroverov, V. N.; Kobayashi, R.; Normand, J.; Raghavachari, K.; Rendell, A.; Burant, J. C.; Iyengar, S. S.; Tomasi, J.; Cossi, M.; Rega, N.; Millam, J. M.; Klene, M.; Knox, J. E.; Cross, J. B.; Bakken, V.; Adamo, C.; Jaramillo, J.; Gomperts, R.; Stratmann, R. E.; Yazyev, O.; Austin, A. J.; Cammi, R.; Pomelli, C.; Ochterski, J. W.; Martin, R. L.; Morokuma, K.; Zakrzewski, V. G.; Voth, G. A.; Salvador, P.; Dannenberg, J. J.; Dapprich, S.; Daniels, A. D.; Farkas, Ö.; Foresman, J. B.; Ortiz, J. V.; Cioslowski, J.; Fox, D. J., Gaussian 09, Revision E.01, Gaussian, Inc., Wallingford CT. 2009.

38. Olsson, M. H. M.; Søndergaard, C. R.; Rostkowski, M.; Jensen, J. H., PROPKA3: Consistent Treatment of Internal and Surface Residues in Empirical pKa Predictions. J. Chem. Theory Comput. 2011, 7 (2), 525–537.

39. Søndergaard, C. R.; Olsson, M. H. M.; Rostkowski, M.; Jensen, J. H., Improved Treatment of Ligands and Coupling Effects in Empirical Calculation and Rationalization of pKa Values. J. Chem. Theory Comput. 2011, 7 (7), 2284–2295.

40. Maier, J. A.; Martinez, C.; Kasavajhala, K.; Wickstrom, L.; Hauser, K. E.; Simmerling, C., ff14SB: Improving the Accuracy of Protein Side Chain and Backbone Parameters from ff99SB. J. Chem. Theory Comput. 2015, 11 (8), 3696–3713.

41. Jorgensen, W. L.; Chandrasekhar, J.; Madura, J. D.; Impey, R. W.; Klein, M. L., Comparison of simple potential functions for simulating liquid water. J. Chem. Phys. 1983, 79 (2), 926–935.

42. Joung, I. S.; Cheatham, T. E., Determination of Alkali and Halide Monovalent Ion Parameters for Use in Explicitly Solvated Biomolecular Simulations. J. Phys. Chem. B 2008, 112 (30), 9020–9041.

43. Marion, A.; Groll, M.; Scharf, D. H.; Scherlach, K.; Glaser, M.; Sievers, H.; Schuster, M.; Hertweck, C.; Brakhage, A. A.; Antes, I.; Huber, E. M., Gliotoxin Biosynthesis: Structure, Mechanism, and Metal Promiscuity of Carboxypeptidase GliJ. ACS Chem. Biol. 2017, 12 (7), 1874–1882.

44. Shannon, R. D.; Fischer, R. X., Empirical electronic polarizabilities in oxides, hydroxides, oxyfluorides, and oxychlorides. Physical Review B 2006, 73 (23), 235111.

45. Loncharich, R. J.; Brooks, B. R.; Pastor, R. W., Langevin dynamics of peptides: The frictional dependence of isomerization rates of N-acetylalanyl-N′-methylamide. Biopolymers 1992, 32 (5), 523–535.

46. Berendsen, H. J.; Postma, J. v.; van Gunsteren, W. F.; DiNola, A.; Haak, J., Molecular dynamics with coupling to an external bath. J. Chem. Phys. 1984, 81 (8), 3684–3690.

47. Ryckaert, J.-P.; Ciccotti, G.; Berendsen, H. J., Numerical integration of the cartesian equations of motion of a system with constraints: molecular dynamics of n-alkanes. J. Comput. Phys. 1977, 23 (3), 327–341.

48. Essmann, U.; Perera, L.; Berkowitz, M. L.; Darden, T.; Lee, H.; Pedersen, L. G., A smooth particle mesh Ewald method. J. Chem. Phys. 1995, 103 (19), 8577–8593.

49. Roe, D. R.; Cheatham, T. E., PTRAJ and CPPTRAJ: Software for Processing and Analysis of Molecular Dynamics Trajectory Data. J. Chem. Theory Comput. 2013, 9 (7), 3084–3095.

50. Andreini, C.; Cavallaro, G.; Lorenzini, S., FindGeo: a tool for determining metal coordination geometry. Bioinformatics 2012, 28 (12), 1658–1660.

51. Liang, Z.; Xue, Y.; Behravan, G.; Jonsson, B. H.; Lindskog, S., Importance of the conserved active‐site residues Try7, Glu106 and Thr199 for the catalytic function of human carbonic anhydrase II. Eur. J. Biochem. 1993, 211 (3), 821–827.

52. Krishnamurthy, V. M.; Kaufman, G. K.; Urbach, A. R.; Gitlin, I.; Gudiksen, K. L.; Weibel, D. B.; Whitesides, G. M., Carbonic Anhydrase as a Model for Biophysical and Physical-Organic Studies of Proteins and Protein−Ligand Binding. Chem. Rev. 2008, 108 (3), 946–1051.

53. Kovalevsky, A.; Aggarwal, M.; Velazquez, H.; Cuneo, M. J.; Blakeley, M. P.; Weiss, K. L.; Smith, J. C.; Fisher, S. Z.; McKenna, R., “To Be or Not to Be” Protonated: Atomic Details of Human Carbonic Anhydrase-Clinical Drug Complexes by Neutron Crystallography and Simulation. Structure 2018, 26 (3), 383–390.e3.

54. Salsbury, F. R.; Crowder, M. W.; Kingsmore, S. F.; Huntley, J. J. A., Molecular dynamic simulations of the metallo-beta-lactamase from Bacteroides fragilis in the presence and absence of a tight-binding inhibitor. J. Mol. Model. 2008, 15 (2), 133.

55. Aitha, M.; Marts, A. R.; Bergstrom, A.; Møller, A. J.; Moritz, L.; Turner, L.; Nix, J. C.; Bonomo, R. A.; Page, R. C.; Tierney, D. L.; Crowder, M. W., Biochemical, Mechanistic, and Spectroscopic Characterization of Metallo-β-lactamase VIM-2. Biochemistry 2014, 53 (46), 7321–7331.

56. Skagseth, S.; Akhter, S.; Paulsen, M. H.; Muhammad, Z.; Lauksund, S.; Samuelsen, Ø.; Leiros, H.-K. S.; Bayer, A., Metallo-β-lactamase inhibitors by bioisosteric replacement: Preparation, activity and binding. Eur. J. Med. Chem. 2017, 135, 159–173.

57. Panteva, M. T.; Giambaşu, G. M.; York, D. M., Comparison of structural, thermodynamic, kinetic and mass transport properties of Mg2+ ion models commonly used in biomolecular simulations. J. Comput. Chem. 2015, 36 (13), 970–982.

58. Zuo, Z.; Liu, J., Assessing the Performance of the Nonbonded Mg2+ Models in a Two-Metal-Dependent Ribonuclease. J. Chem. Inf. Model. 2019, 59 (1), 399–408.

59. Marcos, E. S.; Martínez, J. M.; Pappalardo, R. R., Examining the influence of the [Zn(H2O)6]2+ geometry change on the Monte Carlo simulations of Zn2+ in water. J. Chem. Phys. 1996, 105 (14), 5968–5970.

60. Powell, D. H.; Gullidge, P. M. N.; Neilson, G. W.; Bellissent-Funel, M. C., Zn2+ hydration and complexation in aqueous electrolyte solutions. Mol. Phys. 1990, 71 (5), 1107–1116.

61. MacDermott-Opeskin, H.; McDevitt, C. A.; O’Mara, M. L., Comparing Nonbonded Metal Ion Models in the Divalent Cation Binding Protein PsaA. J. Chem. Theory Comput. 2020, 16 (3), 1913–1923.

