## Supporting Information for "Assessment of Molecular Mechanics-based Zn^2+^ Models in Mono- and Bimetallic Ligand Binding Sites"

|

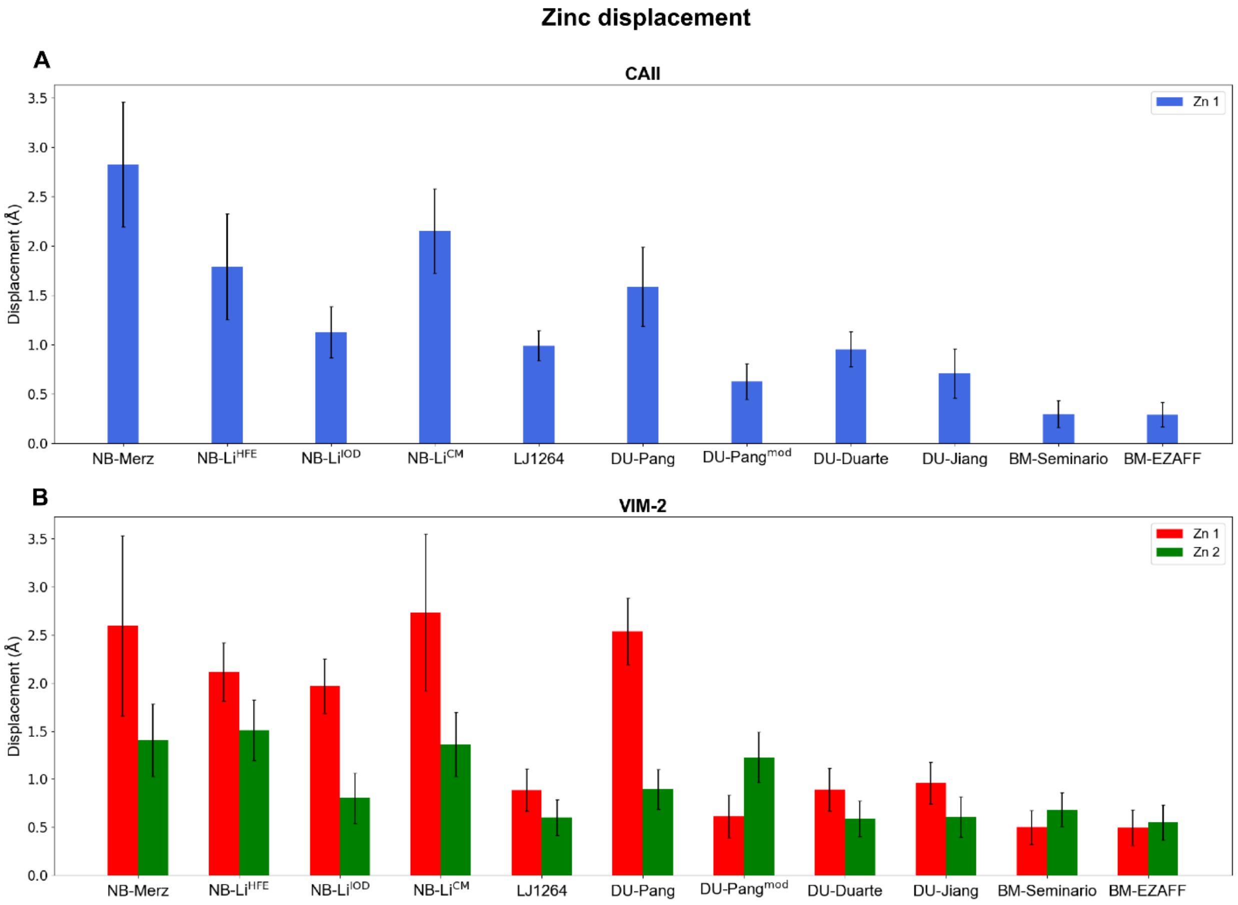

**Figure S1.** Zinc displacement observed in CAII (A) and VIM-2 (B). The displacement is calculated with respect to the position of the $Zn^{2+}$ ion in the X-Ray structure.

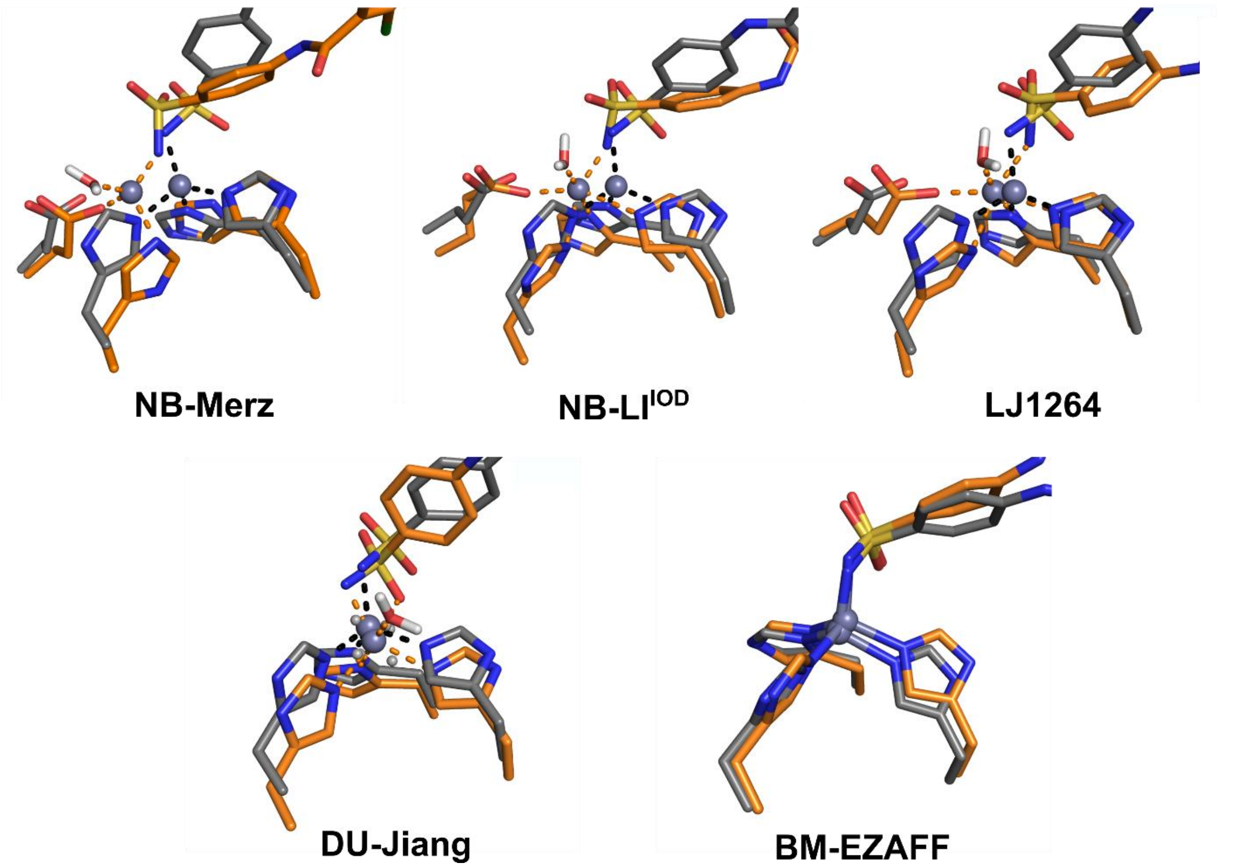

**Figure S2.** Sampled coordination geometries by selected $Zn^{2+}$ models in CAII. Carbon atoms of the reference structure and the simulated conformations are shown in grey and orange, respectively. Dotted lines represent the coordination geometry.

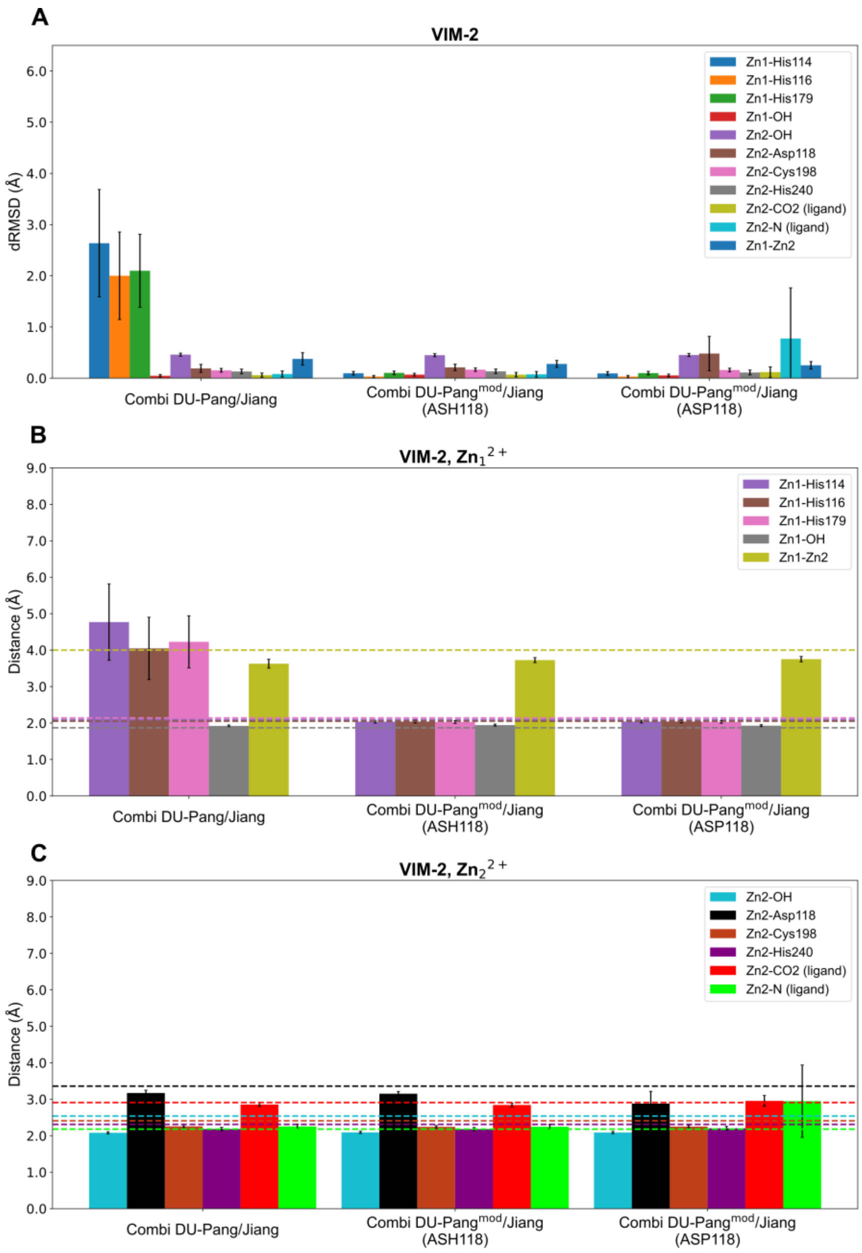

**Figure S3.** Performance of combining DU-Pang and DU-Jiang for the bimetallic system VIM-2. DU-Pang was applied for $Zn_1^{2+}$, DU-Jiang for $Zn_2^{2+}$. Simulations were performed with DU-Pang or DU-Pang$^{mod}$ (as described in the main text), with Asp118 either protonated (ASH118) or deprotonated (ASP118). A: RMSD of interatomic distances (dRMSD) between $Zn^{2+}$ and selected binding site atoms with respect to the average simulated interatomic distance of the respective atom pair as a measure for the stability of the sampled $Zn^{2+}$ binding site. B&C: Interatomic distance between $Zn_1^{2+}$ (B) or $Zn_2^{2+}$ (C) and selected binding site atoms, as a measure for binding site integrity. The bars represent the average distance between the $Zn^{2+}$ ion and the ligating residue during the simulation, while the dotted lines represent the value of the respective distance in the X-Ray structure. All distances to an aspartate or glutamate residue are measured to the side-chain's carboxyl carbon, since both oxygen atoms are equivalent and can freely take over each-other's function.

**Table S1.** Bonded Parameters Applied in Bonded Models

| New atomtype | Applied on |
|---|---|
| *CAII* | |
| **M1** | Zn1 |
| **Y1** | His94 (ND) |
| **Y2** | His96 (ND) |
| **Y3** | His119 (NE) |
| **Y4** | Lig (N) |
| *VIM-2* | |
| **M1** | Zn1 |
| **M2** | Zn2 |
| **Y1** | His114 (NE) |
| **Y2** | His116 (ND) |
| **Y3** | His179 (NE) |
| **Y4** | OH- (O) |
| **Y5** | Asp118 (OD2) |
| **Y6** | Cys198 (SG) |
| **Y7** | His240 (NE) |
| **Y8** | Ligand (O) |
| **Y9** | Ligand (N) |

| ***bonds*** | **BM-Seminario** | | **BM-EZAFF** | |
|---|---|---|---|---|
| | **$R_{eq}$ (Å)** | **Fc (kcal·mol$^{-1}$·A$^{-2}$)** | **$R_{eq}$ (Å)** | **Fc (kcal·mol$^{-1}$·A$^{-2}$)** |
| *CAII* | | | | |
| **M1-Y1** | 2.0316 | 68.8 | 2.049 | 62.2 |
| **M1-Y2** | 2.0531 | 60.6 | 2.092 | 48.4 |
| **M1-Y3** | 2.0314 | 68.4 | 2.061 | 58.1 |
| **M1-Y4** | 1.9615 | 100.1 | 1.993 | 85.8 |
| *VIM-2* | | | | |
| **M1-Y4** | 1.8763 | 132.9 | 1.875 | 154 |
| **M2-Y4** | 2.1105 | 32.7 | 2.541 | 16.4 |
| **M2-Y8** | 2.1924 | 22.6 | 2.201 | 29.5 |
| **M2-Y9** | 2.5153 | 0.00 | 2.175 | 29 |
| **Y1-M1** | 2.0561 | 58.3 | 2.133 | 37.8 |
| **Y2-M1** | 2.0993 | 44.3 | 2.047 | 63 |
| **Y3-M1** | 2.0616 | 56.1 | 2.129 | 38.5 |
| **Y5-M2** | 2.1175 | 12.2 | 2.324 | 21.2 |
| **Y6-M2** | 2.3917 | 41.1 | 2.41 | 36.3 |
| **Y7-M2** | 2.2002 | 26.3 | 2.312 | 11.3 |

| *angles* | BM-Seminario | | BM-EZAFF | |
|---|---|---|---|---|
| | **$\theta_{eq}$** | **Fc (kcal·mol$^{-1}$·rad$^{-2}$)** | **$\theta_{eq}$** | **Fc (kcal·mol$^{-1}$·rad$^{-2}$)** |
| *CAII* | | | | |
| **Y1-M1-Y2** | 105.76 | 32.42 | 104.49 | 35 |
| **Y1-M1-Y3** | 107.57 | 31.84 | 111.69 | 35 |
| **Y1-M1-Y4** | 116.37 | 30.26 | 111.23 | 35 |
| **Y2-M1-Y3** | 110.86 | 38.74 | 98.57 | 35 |
| **Y2-M1-Y4** | 100.97 | 49.16 | 112.38 | 35 |
| **Y3-M1-Y4** | 114.76 | 50.63 | 117.14 | 35 |
| **CC-Y3-M1** | 133.2 | 54.85 | 130.73 | 50 |
| **CR-Y1-M1** | 123.55 | 50.7 | 128.77 | 50 |
| **CR-Y2-M1** | 124.61 | 53.78 | 129.46 | 50 |
| **M1-Y1-CV** | 129.89 | 54.18 | 122.39 | 50 |
| **M1-Y2-CV** | 128.85 | 52.21 | 121.46 | 50 |
| **M1-Y3-CR** | 119.58 | 52.85 | 120.65 | 50 |
| **M1-Y4-hn** | 116.69 | 45.25 | 120 | 50 |
| **M1-Y4-sy** | 115.75 | 73.09 | 115.3 | 50 |
| *VIM-2* | | | | |
| **Y1-M1-Y2** | 103.38 | 31.1 | 100.44 | 35 |
| **Y1-M1-Y3** | 106.23 | 33.22 | 104.34 | 35 |
| **Y1-M1-Y4** | 118.17 | 35.38 | 109.85 | 35 |
| **Y2-M1-Y3** | 103.64 | 36.16 | 107.64 | 35 |
| **Y2-M1-Y4** | 107.75 | 46.34 | 117.18 | 35 |
| **Y3-M1-Y4** | 116 | 45.97 | 115.61 | 35 |
| **Y4-M2-Y8** | 85.24 | 45.75 | 95.05 | 35 |
| **Y4-M2-Y9** | 79.04 | 17.24 | 82.98 | 35 |
| **Y5-M2-Y4** | 93.53 | 71.49 | 76.41 | 35 |
| **Y5-M2-Y6** | 99.46 | 13.47 | 100.91 | 35 |
| **Y5-M2-Y7** | 86.44 | 37.55 | 92.92 | 35 |
| **Y5-M2-Y8** | 155.6 | 21.26 | 168.77 | 35 |
| **Y5-M2-Y9** | 83.85 | 9.96 | 91.91 | 35 |
| **Y6-M2-Y4** | 102.6 | 26.95 | 95.01 | 35 |
| **Y6-M2-Y7** | 95.07 | 19.09 | 93.58 | 35 |
| **Y6-M2-Y8** | 104.6 | 31.08 | 86.94 | 35 |
| **Y6-M2-Y9** | 176.16 | 14.56 | 166.21 | 35 |
| **Y7-M2-Y4** | 162.08 | 29.07 | 167.4 | 35 |
| **Y7-M2-Y8** | 87.44 | 29.25 | 94.6 | 35 |
| **Y7-M2-Y9** | 83.14 | 14.26 | 90.8 | 35 |
| **Y9-M2-Y8** | 71.97 | 19.08 | 79.67 | 35 |
| **CC-Y2-M1** | 127.25 | 61.04 | 121.61 | 50 |
| **CO-Y5-M2** | 128.33 | 20.92 | 142.67 | 50 |
| **CR-Y1-M1** | 120.44 | 49.09 | 126.33 | 50 |
| **CR-Y3-M1** | 122.37 | 53.68 | 133.62 | 50 |
| **CR-Y7-M2** | 124.06 | 44.83 | 120.28 | 50 |

*Continues on next page…*

| | | | | |
|---|---|---|---|---|
| **CT-Y6-M2** | **103.02** | **66.62** | **114.99** | **70** |
| **M1-Y1-CV** | 132.34 | 51.82 | 122.52 | 50 |
| **M1-Y2-CR** | 123.84 | 63.83 | 129.15 | 50 |
| **M1-Y3-CV** | 130.26 | 61.87 | 115.25 | 50 |
| **M1-Y4-M2** | 133.89 | 40.22 | 129.5 | 50 |
| **M2-Y7-CV** | 128.71 | 40.02 | 129.61 | 50 |
| **M2-Y8-c** | 120.04 | 37.35 | 109.99 | 50 |
| **M2-Y9-cc** | 105.06 | 17.82 | 105.39 | 50 |
| **M2-Y9-cd** | 141.12 | 16.49 | 134.59 | 50 |
| **M1-Y4-HW** | 111.06 | 32.46 | 120 | 50 |
| **M2-Y4-HW** | 96.26 | 31.62 | 120 | 50 |

Torsion barriers in newly introduced dihedrals are by default set to 0 in this method.

**Table S2.** Overview of Heat-up Protocol Applied in this Study.

| Step[a] | Time (ps) | Final temperature (K) | Ensemble | Weight restraint (kcal·mol·Å$^{-2}$) | Restrained atoms[b] |
|---|---|---|---|---|---|
| H-1 | 20 | 0 | NVT | 20 | All atoms |
| H-2 | 120 | 5 | NVT | 20 | All atoms |
| H-3 | 220 | 10 | NVT | 20 | All atoms |
| H-4 | 320 | 20 | NVT | 20 | All atoms |
| H-5 | 420 | 50 | NVT | 20 | BB+BiS |
| H-6 | 620 | 100 | NVT | 20 | BB+BiS |
| Eq-7 | 820 | 100 | NVT | 20 | BB+BiS |
| H-8 | 1220 | 200 | NVT | 20 | BB+BiS |
| Eq-9 | 2220 | 200 | NVT | 20 | BiS |
| H-10 | 3220 | 300 | NVT | 20 | BiS |
| Eq-11 | 4220 | 300 | NPT | 20 | BiS |
| Eq-12 | 4720 | 300 | NPT | 15 | BiS |
| Eq-13 | 5220 | 300 | NPT | 10 | BiS |
| Eq-14 | 5720 | 300 | NPT | 5 | BiS |
| Eq-15 | 6220 | 300 | NPT | 4 | BiS |
| Eq-16 | 6720 | 300 | NPT | 3 | BiS |
| Eq-17 | 7220 | 300 | NPT | 2 | BiS |
| Eq-18 | 7720 | 300 | NPT | 1 | BiS |
| Eq-19 | 8720 | 300 | NPT | 0 | - |

a. H: heat-up, Eq: equilibration
b. BB: backbone atoms (Cα,C,N), BiS: binding site, consisting of all residues with at least one atom within a 5 Å sphere of the $Zn^{2+}$ ion(s).

**Table S3.** Distribution of $Zn^{2+}$ Coordination Geometries, Covering All Coordination Geometries, in Percentage. CAII.

| **Coordination geometry** | NB-Merz | NB-$Li^{HFE}$ | NB-$Li^{IOD}$ | NB-$Li^{CM}$ | LJ1264 | DU-Pang | DU-$Pang^{mod}$ | DU-Duarte | DU-Jiang | BM-Seminaro | BM-EZAFF |
|---|---|---|---|---|---|---|---|---|---|---|---|
| Tetrahedral[a] | 1.33 | 99.33 | 0.00 | 0.67 | 0.00 | 20.67 | 96.67 | 0.00 | 0.00 | 80.67 | 84.00 |
| Octahedral | 86.67 | 0.00 | 94.67 | 96.00 | 95.33 | 18.67 | 0.00 | 98.00 | 82.67 | 0.00 | 0.00 |
| Octahedral (capped face) | 1.33 | 0.00 | 0.00 | 0.00 | 0.00 | 0.67 | 0.00 | 0.00 | 12.00 | 0.00 | 0.00 |
| Square pyramidal | 4.00 | 0.00 | 1.33 | 1.33 | 0.00 | 4.00 | 0.00 | 0.00 | 0.00 | 0.00 | 0.00 |
| Pentagon bipyramidal + vac | 0.00 | 0.00 | 3.33 | 0.00 | 4.67 | 4.67 | 0.00 | 2.00 | 5.33 | 0.00 | 0.00 |
| Trigonal prism | 0.00 | 0.00 | 0.00 | 0.00 | 0.00 | 0.67 | 0.00 | 0.00 | 0.00 | 0.00 | 0.00 |
| Trigonal prism + vac | 0.00 | 0.00 | 0.00 | 0.00 | 0.00 | 2.00 | 0.00 | 0.00 | 0.00 | 0.00 | 0.00 |
| Trigonal bipyramidal | 0.00 | 0.00 | 0.00 | 2.00 | 0.00 | 10.00 | 1.33 | 0.00 | 0.00 | 0.00 | 0.00 |
| Trigonal bipyramidal + vac | 0.00 | 0.00 | 0.00 | 0.00 | 0.00 | 0.67 | 0.00 | 0.00 | 0.00 | 0.00 | 0.00 |
| Irregular | 6.67 | 0.67 | 0.67 | 0.00 | 0.00 | 38.00 | 2.00 | 0.00 | 0.00 | 19.33 | 16.00 |

[a] Native coordination geometry, as observed in the X-Ray structure for CAII (PDB ID: 5NXG).
vac = vacancy

**Table S4.** Distribution of $Zn^{2+}$ Coordination Geometries, Covering All Coordination Geometries, in Percentage. VIM-2.

| **Coordination geometry** | NB-Merz | NB-$Li^{HFE}$ | NB-$Li^{IOD}$ | NB-$Li^{CM}$ | LJ1264 | DU-Pang | DU-$Pang^{mod}$ | DU-Duarte | DU-Jiang | DU-com1 | Du-com2 | DU-com3 | BM-Seminaro | BM-EZAFF |
|---|---|---|---|---|---|---|---|---|---|---|---|---|---|---|
| | | | | | | | $Zn_1^{2+}$ geometry occupancy (%) | | | | | | | |
| Tetrahedral[a] | 0.00 | 96.00 | 0.00 | 0.00 | 0.00 | 4.67 | 99.33 | 0.00 | 0.00 | 20.67 | 99.33 | 100.00 | 89.33 | 94.00 |
| Octahedral | 95.33 | 0.00 | 100.00 | 96.67 | 100.00 | 22.67 | 0.00 | 100.00 | 100.0 | 6.67 | 0.00 | 0.00 | 0.00 | 0.00 |
| Octahedral (capped face) | 0.00 | 0.00 | 0.00 | 0.00 | 0.00 | 1.33 | 0.00 | 0.00 | 0.00 | 0.67 | 0.00 | 0.00 | 0.00 | 0.00 |
| Octahedral (monocapped) + vac | 0.00 | 0.00 | 0.00 | 0.00 | 0.00 | 0.00 | 0.00 | 0.00 | 0.00 | 0.00 | 0.00 | 0.00 | 0.00 | 0.00 |
| Square pyramidal | 4.00 | 0.00 | 0.00 | 2.67 | 0.00 | 0.00 | 0.00 | 0.00 | 0.00 | 0.00 | 0.00 | 0.00 | 0.00 | 0.00 |
| Trigonal bipyramidal | 0.00 | 1.33 | 0.00 | 0.00 | 0.00 | 40.67 | 0.00 | 0.00 | 0.00 | 41.33 | 0.00 | 0.00 | 0.00 | 0.67 |
| Trigonal bipyramidal + vac | 0.00 | 0.00 | 0.00 | 0.00 | 0.00 | 0.00 | 0.67 | 0.00 | 0.00 | 2.00 | 0.00 | 0.00 | 10.00 | 0.67 |
| Pentagonal bipyramidal + vac | 0.00 | 0.00 | 0.00 | 0.00 | 0.00 | 3.33 | 0.00 | 0.00 | 0.00 | 0.00 | 0.00 | 0.00 | 0.00 | 0.00 |
| Irregular | 0.67 | 2.67 | 0.00 | 0.67 | 0.00 | 27.33 | 0.00 | 0.00 | 0.00 | 28.67 | 0.67 | 0.00 | 0.67 | 4.67 |
| | | | | | | | $Zn_2^{2+}$ geometry occupancy (%) | | | | | | | |
| Tetrahedral | 98.00 | 100.00 | 0.00 | 89.33 | 0.00 | 22.00 | 92.00 | 0.00 | 0.00 | 0.00 | 0.00 | 0.00 | 0.00 | 0.00 |
| Octahedral[a] | 0.00 | 0.00 | 43.33 | 0.00 | 99.33 | 0.67 | 0.00 | 100.00 | 100.00 | 100.00 | 100.00 | 92.67 | 100.00 | 100.00 |
| Octahedral (capped face) | 0.00 | 0.00 | 0.67 | 0.00 | 0.00 | 0.00 | 0.00 | 0.00 | 0.00 | 0.00 | 0.00 | 0.00 | 0.00 | 0.00 |
| Square pyramidal | 0.00 | 0.00 | 0.00 | 0.00 | 0.00 | 0.00 | 0.00 | 0.00 | 0.00 | 0.00 | 0.00 | 0.00 | 0.00 | 0.00 |
| Trigonal bipyramidal | 0.67 | 0.00 | 0.00 | 2.00 | 0.00 | 17.33 | 0.00 | 0.00 | 0.00 | 0.00 | 0.00 | 0.00 | 0.00 | 0.00 |
| Pentagonal bipyramidal + vac | 0.00 | 0.00 | 37.33 | 0.00 | 0.67 | 0.67 | 0.00 | 0.00 | 0.00 | 0.00 | 0.00 | 7.33 | 0.00 | 0.00 |
| Trigonal prism + vac | 0.00 | 0.00 | 0.00 | 0.00 | 0.00 | 1.34 | 0.00 | 0.00 | 0.00 | 0.00 | 0.00 | 0.00 | 0.00 | 0.00 |
| Irregular | 1.33 | 0.00 | 18.67 | 8.67 | 0.00 | 58.00 | 8.00 | 0.00 | 0.00 | 0.00 | 0.00 | 0.00 | 0.00 | 0.00 |

[a] Native coordination geometry, as observed in the X-Ray structure for VIM-2 (PDB ID: 6HF5).
DU-com1: Combined DU-Pang/DU-Jiang model for $Zn_1^{2+}$ and $Zn_2^{2+}$ respectively.
DU-com2: Combined DU-$Pang^{mod}$/DU-Jiang model for $Zn_1^{2+}$ and $Zn_2^{2+}$ respectively.
DU-com3: Combined DU-$Pang^{mod}$/DU-Jiang model for $Zn_1^{2+}$ and $Zn_2^{2+}$ respectively, with Asp118 deprotonated.
vac = vacancy
